# Multi-domain O-GlcNAcase structures reveal allosteric regulatory mechanisms

**DOI:** 10.1101/2025.03.10.642372

**Authors:** Sara Basse Hansen, Sergio G. Bartual, Huijie Yuan, Olawale G. Raimi, Andrii Gorelik, Andrew T. Ferenbach, Kristian Lytje, Jan Skov Pedersen, Taner Drace, Thomas Boesen, Daan M. F. van Aalten

## Abstract

Nucleocytoplasmic protein O-GlcNAcylation is an essential modification catalysed by *O*-GlcNAc transferase (OGT) and reversed by O-GlcNAc hydrolase (OGA), a multi-domain enzyme that also contains a C-terminal pseudo-histone acetyltransferase (pHAT) domain. OGA and OGT are tightly regulated using O-GlcNAc-dependent feedback mechanisms that are largely unknown. Although the structure of the OGA homodimeric catalytic domain has been reported, the structure and function of the pHAT domain remains poorly understood. We describe a crystal structure of the *Trichoplax adhaerens* pHAT domain and cryo-EM data of the multi-domain *T. adhaerens* and human OGAs, together with surface plasmon resonance and small-angle X-ray scattering studies. Our findings show that the eukaryotic OGA pHAT domains affect O-GlcNAc homeostasis, and form catalytically incompetent, symmetric homodimers, projecting a partially conserved putative peptide binding site available for interactions with binding partners. While there is evidence for symmetric OGA multi-domain dimers in solution, interactions between the linkers to the pHAT domains allow these to adopt a limited range of positions. In hOGA, the positions of the pHAT domains determine the wider active site environment through a key conformational change involving a tryptophan in a flexible arm region. Taken together, these multi-domain OGA structures reveal allosteric mechanisms of regulation.

## Introduction

Nucleocytoplasmic modification of 1000s of proteins with O-linked N-acetylglucosamine (O-GlcNAc) is a dynamic process (Torres & Hart, 1984; Haltiwanger *et al*., 1992; Wulff-Fuentes *et al*., 2021) and modulates a broad range of critical cellular processes such as metabolism, translation, stress response, transcription and protein homeostasis (Yang & Qian, 2017; Wulff-Fuentes *et al*., 2021). O-GlcNAcylation is mediated by a single pair of enzymes: the *O*-GlcNAc transferase (OGT) that adds O-GlcNAc to serine and threonine residues, and the O-GlcNAcase (OGA), which removes *O*-GlcNAc from the modified proteins (Yang & Qian, 2017; Haltiwanger *et al*., 1992; Dong & Hart, 1994). This modification is particularly critical for development of the central nervous system in vertebrates, and *de novo* missense variants in *OGT* and *OGA* have been associated with neurodevelopmental disorders (Pravata *et al*., 2020; Mayfield *et al*., 2024; Authier *et al*., 2023). In metazoans the O-GlcNAc system is conserved across all taxa (Shafi *et al*., 2000), including the simplest multicellular animal known to possess a functional O-GlcNAc system, the metazoan *Trichoplax adhaerens* (Selvan *et al*., 2015). *T. adhaerens* possesses two *OGA* genes producing two functional proteins, a shorter *Ta*OGA53, lacking the C-terminal domain, and a longer *Ta*OGA54 (the latter referred to as *Ta*OGA throughout this manuscript) (Selvan *et al*., 2015), whereas vertebrates possess several OGA isoforms produced by alternative splicing of a single *OGA* gene (also known as “meningioma expressed antigen 5” *(mgea5*) (Comtesse *et al*., 2001)).

The human full-length OGA (hOGA), considered to be the primary isoform, is found in the nucleocytoplasmic space as an obligate dimer and is composed of three domains **(Fig. 1A)**. The N-terminal catalytic domain belongs to the CAZy GH84 glycosyl hydrolase family and employs a substrate-assisted catalytic mechanism to hydrolyse O-GlcNAc from target proteins (Dennis *et al*., 2006; Rao *et al*., 2006; Macauley *et al*., 2005) This active site is a target for competitive inhibitors designed to increase O-GlcNAc levels, offering possible therapeutic potential for neurodevelopmental diseases such as Alzheimer’s and Parkinson disease (Bartolomé-Nebreda *et al*., 2021; Selnick *et al*., 2019). Recent crystal structures of a truncated hOGA excluding the C-terminal domain reveal a homodimer where the helical stalk domain **(Fig. 1A)** contributes to the dimer interface (Li, Li, Lu *et al*., 2017; Roth *et al*., 2017; Li, Li, Hu *et al*., 2017). Furthermore, a cryo-EM structure of the hOGA catalytic core in complex with human OGT was recently reported (Lu *et al*., 2023). Together with earlier studies of substrate complexes with a bacterial OGA orthologue (Schimpl *et al*., 2010), these hOGA structures reveal that substrate recognition is shaped by regions beyond the active site. For instance, the stalk domain, which consists of a helical bundle and an ‘arm region’ (residues 665-685) close to the active site, plays a crucial role in shaping substrate accessibility by contributing to formation of the peptide-binding cleft that is capable of binding glycopeptides in a bidirectional mode (Li, Li, Hu *et al*., 2017). Mutations in the stalk domain can affect hOGA activity and have been implicated in cancer progression (Hu *et al*., 2023). For instance, mutations such as S652F and R586A disrupt OGA-substrate interactions (Hu *et al*., 2023). Specifically, the S652F mutation enhances O-GlcNAc removal from PDLIM7, ultimately suppressing p53 gene expression and promoting malignancy (Hu *et al*., 2023).

**Fig. 1:**
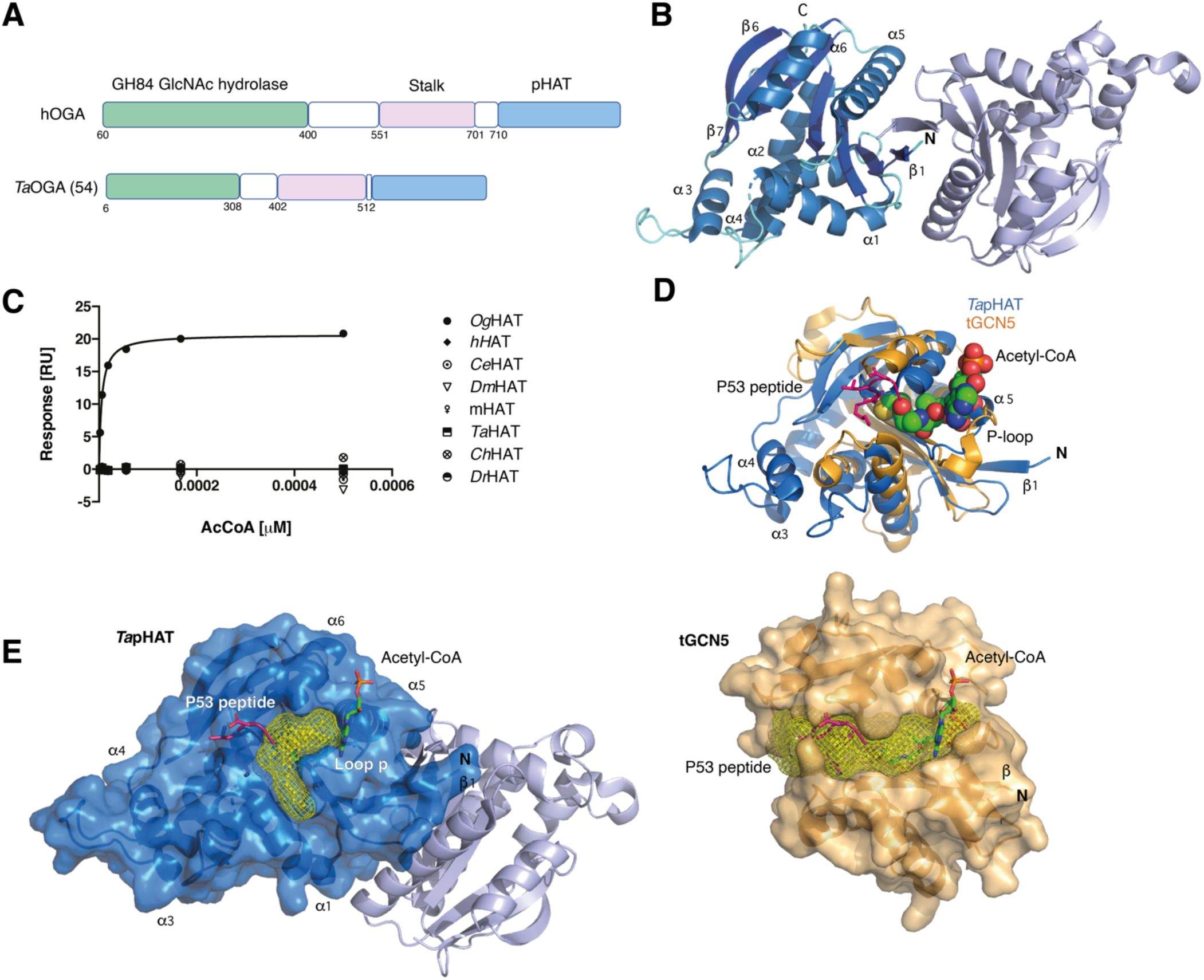
Structure and conservation of the *Ta*pHAT domain dimer. **A)** Schematic representation of hOGA and *Ta*OGA domain organisation. The white sections represent regions of predicted disorder. **B)** Cartoon representation of *Ta*pHAT domain structure. Monomer A α-helices are coloured in marine blue, the β-sheets in blue, while the unstructured regions are coloured aquamarine. Monomer B is coloured in slate. **C)** SPR experiment showing acetyl-CoA (AcCoA) binding by *Og*AT (positive control) but not by metazoan proteins (h – human, Ce – *C. elegans*, Dm – *D. melanogaster*, m – mouse, Ta – *T. adhaerens*, Ch – chicken, Dr – *D. rerio*). **D)** Superposition of the *Ta*pHAT domain (marine) with the tGCN5 structure (PDB 1Q2D (Poux & Marmorstein, 2003), bright orange) co-crystallised with acetyl-CoA (spheres) and a P53 peptide (hot pink). *Ta*pHAT domain structural elements are annotated for easy reference. **E)** *Ta*pHAT domain dimer overlayed with the ligands extracted from the tGCN5 structure (left). Only monomer A surface is shown (marine). The surface representation of the tGCN5 structure (bright orange) is represented on the right for comparison. Cavities were detected with the PyMOL plugin CavitOmix (Hetmann *et al*., 2023) and represented as yellow meshes in both structures.

While the OGA C-terminal domain **(Fig. 1A)** is predicted to have a fold similar to the GCN5 histone *N*-acetyltransferase family and was initially reported to have histone acetyltransferase activity *in vitro* (Toleman *et al*., 2004), subsequent studies failed to support this (Rao, Schüttelkopf *et al*., 2013; Butkinaree *et al*., 2008), and the domain is now regarded to be a non-catalytic pseudo-histone acetyltransferase (pHAT) domain of unknown function. Nevertheless, mutational evidence implicates the pHAT domain in (regulation of) acetylation processes. For instance, the Y891F mutation in the pHAT domain has been reported to impair the acetylation of PKM2 in cells, yet purified OGA does not directly acetylate PKM2, leading to the hypothesis that the pHAT domain acts as a scaffold, recruiting an unknown acetyltransferase (Singh *et al*., 2020). This suggests that the pHAT domain may have evolved to mediate specific protein-protein interactions and/or allosteric regulation to OGA rather than retain catalytic function. Interestingly, a missense *OGA* variant in the pHAT domain has recently been associated with neurodevelopmental delay (Authier *et al*., 2023) further highlighting the significance of this domain. *OGT* missense variants leading to intellectual disability are associated with a (compensatory) loss of OGA mRNA and protein, leading to the intriguing possibility that the resulting loss of the functions associated with the pHAT domain could also contribute to the mechanisms underpinning this disease.

The pHAT domain substantially influences OGA activity. OGA lacking the pHAT domain exhibits a 20-fold lower *K*_m_ for certain substrates compared to the full-length protein, while other substrates were restricted from modification (Forsythe *et al*., 2006) (Keembiyehetty *et al*., 2011). The pHAT domain has been reported to be involved in translocation of hOGA to the nucleus in response to DNA damage (Liu *et al*., 2021), where it modulates transcription by altering the O-GlcNAcylation patterns on transcription factors, RNA polymerase II and nucleosomes (Liu *et al*., 2024; Lewis, 2024; Dupas *et al*., 2023). However, it is not known how the pHAT domains orchestrate the translocation of hOGA to the nuclear space, or whether they are required for interactions with chromatin-associated proteins.

To maintain O-GlcNAc homeostasis, OGA is known to be subjected to multiple layers of regulation. Transcriptional and translational feedback mechanisms adjust *OGA* expression in minutes to hours in response to cellular needs (Zhang *et al*., 2014). However, more immediate regulation on shorter time scales could also be achieved through post-translational modifications such as Ser406 O-GlcNAcylation (Gorelik *et al*., 2019) and/or allosteric effects that could involve the non-catalytic domains of OGA, such as the pHAT domains (Huang *et al*., 2020). In summary, it is still not understood what the functions of the pHAT domains are, how these domains are positioned relative to the OGA homodimer catalytic cores and how they contribute to regulating OGA activity.

Here, we report the crystal structure of the isolated pHAT domain homodimer and the cryo-EM full-length homodimeric structure of *Trichoplax adhaerens* OGA (*Ta*OGA). Together with SPR experiments, these data show that while the pHAT domains from eukaryotic OGAs regulate O-GlcNAc homeostasis and retain a putative acceptor binding site, they cannot bind acetyl-CoA and are therefore pseudo-histone acetyl transferases. Linkers rich in negative charge appear to decouple the *T. adhaerens* pHAT domains from the symmetric homodimer catalytic core, allowing the pHAT domains to adopt undefined positions. Furthermore, cryo-EM structures of the multi-domain hOGA homodimer reveal that its pHAT domains are tethered via proline-rich flexible linkers that break the two-fold symmetry present in the catalytic core homodimers. This forces hOGA to adopt a limited range of asymmetric conformations with exposed putative peptide binding sites on the pHAT domain, further supported by solution small angle scattering experiments. The positions of the pHAT domains determine the wider active site environment through a key conformational change involving a tryptophan in a flexible arm region. Taken together, these multi-domain OGA structures reveal previously unknown molecular mechanisms of allosteric regulation.

## Results & Discussion

### The pHAT domains of eukaryotic OGAs form homodimers

Although hOGA is a multi-domain protein, including a C-terminal pHAT domain, structural studies of the full-length enzyme have been hampered by the presence of several large regions of predicted disorder **(Fig. 1A)**. Indeed, in three studies describing crystal structures of the homodimeric hOGA catalytic core these regions (and the pHAT domains) were not included in the expression constructs (Roth *et al*., 2017; Li, Li, Lu *et al*., 2017; Li, Li, Hu *et al*., 2017). To facilitate an understanding of the structure and function of the pHAT domain, we turned to the evolutionary ancient and simple organism *Trichoplax adhaerens,* that has been shown to possess a functional OGA with simpler architecture **(Fig. 1A)** (Selvan *et al*., 2015). As a first step towards a full-length *Ta*OGA structure, we determined the crystal structure of the *Ta*OGA pHAT (*Ta*pHAT) domain. Protein produced in *E. coli* produced crystals suitable for structure determination at 1.8 Å (**Fig. 1B**, **Table 1**). The *Ta*pHAT structure reveals two molecules in the asymmetric unit, with their *β*1-strands packing in a parallel fashion **(Fig. 1B)**. Consistent with the predicted pHAT canonical GCN5 acetyltransferase fold (Dyda *et al*., 2000), the overall topology of *Ta*pHAT consists of a central core of seven anti-parallel *β*-sheets surrounded by six *α*-helices (**Fig. 1B**). Furthermore, the 995 Å^2^ interdomain interface surface area, calculated with PISA (Krissinel & Henrick, 2007), together with the *Ta*pHAT chromatography elution profiles (**Supplementary Fig. 1A**) suggest that the dimer observed *in crystallo* is also present in solution. This would be in agreement with the homodimeric nature of the hOGA catalytic core crystal structures published to date (Roth *et al*., 2017; Li, Li, Lu *et al*., 2017; Li, Li, Hu *et al*., 2017). To assess whether this pHAT homodimeric arrangement is observed in other GCN5 family members, we first used the PISA server (Krissinel & Henrick, 2007) to find structures with an interface at least 70% similar to the observed *Ta*pHAT dimeric interface, but none were found. Then, we used PISA to search for proteins with an interface similar to the *Ta*pHAT monomer, regardless of their oligomeric arrangement. Among the hits, only two structures had a Q score (1 for identical, 0 for dissimilar proteins) higher than 0.5, the *Oceanicola granulosus* bacterial HAT (*Og*HAT) (PDB 3ZJ0 (Rao, Schüttelkopf *et al*., 2013), Q score 0.62) and the *Saccharomyces cerevisiae* pseudo-HAT domain (*Ss*pHAT) (PDB 4BMH (He *et al*., 2014a) Q score 0.54), although neither of these are known to form dimers. The closest hit reported to form a biological dimer is the histone acetyltransferase protein from the archaeon *Pyrococcus horikoshii* (*Ph*HAT) (PDB 1WWZ (Kunishima, N, 2005) Q score 0.42), however inspection of the structural alignment between *Ta*pHAT and *Ph*HAT reveals that the homodimer interface is structurally dissimilar **(Supplementary Fig. 1B)**. In *Ta*pHAT the dimer interface is mostly formed by the parallel packing of the *β*1 strands plus the interaction of the *α*5 helix from one monomer with the middle part of the *β*1 strand from the other monomer. In contrast, the *Ph*HAT dimeric interface is exclusively formed by *α*-helical interactions, under this arrangement the *β*1 strands are pushed away from the dimerisation interface (**Supplementary Fig. 1B**).

**Table 1:**
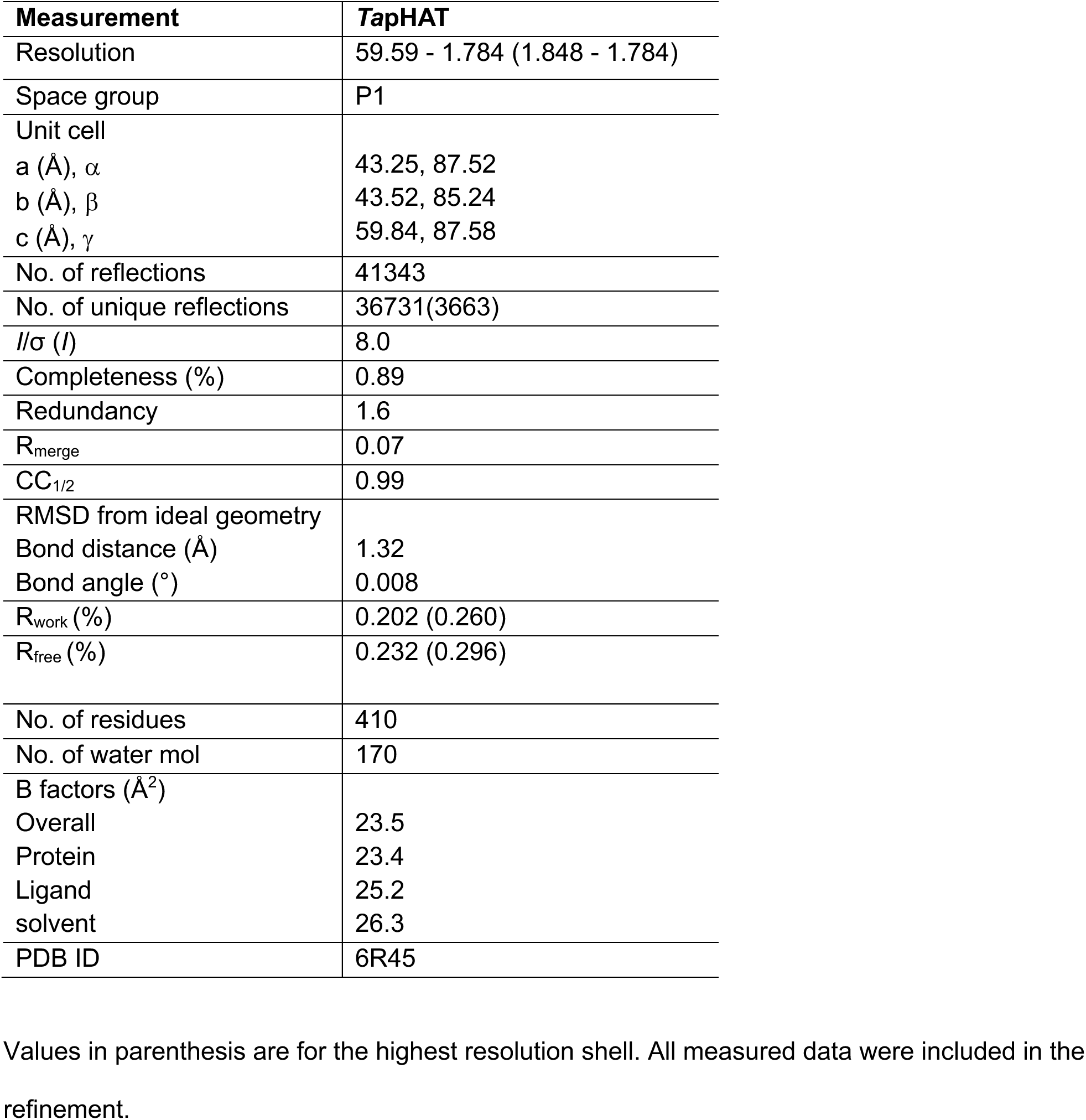
Diffraction data and refinement statistics for the *Ta*pHAT crystal structure.

| Measurement | *TapHAT* |
| --- | --- |
| Resolution | 59.59 - 1.784 (1.848 - 1.784) |
| Space group | P1 |
| Unit cell |  |
| a (Å), $\alpha$ | 43.25, 87.52 |
| b (Å), $\beta$ | 43.52, 85.24 |
| c (Å), $\gamma$ | 59.84, 87.58 |
| No. of reflections | 41343 |
| No. of unique reflections | 36731(3663) |
| $I/\sigma(I)$ | 8.0 |
| Completeness (%) | 0.89 |
| Redundancy | 1.6 |
| $R_{\text{merge}}$ | 0.07 |
| $CC_{1/2}$ | 0.99 |
| RMSD from ideal geometry |  |
| Bond distance (Å) | 1.32 |
| Bond angle (°) | 0.008 |
| $R_{\text{work}}$ (%) | 0.202 (0.260) |
| $R_{\text{free}}$ (%) | 0.232 (0.296) |
| No. of residues | 410 |
| No. of water mol | 170 |
| B factors (Å<sup>2</sup>) |  |
| Overall | 23.5 |
| Protein | 23.4 |
| Ligand | 25.2 |
| solvent | 26.3 |
| PDB ID | 6R45 |

Values in parenthesis are for the highest resolution shell. All measured data were included in the refinement.

We next investigated whether the *Ta*pHAT dimer interface is conserved among the pHAT domains of eukaryotic OGAs. A search of the predicted *Ta*pHAT interface in the PISA server (Krissinel & Henrick, 2007) reveals that, despite the *β*1 strand being conserved across species (**Supplementary Figs. 1C,2**), a network of electrostatic interactions in the *Ta*pHAT dimer is not present in other eukaryotic pHAT domains **(Supplementary Figs. 1C,2)** – perhaps explaining why other pHAT domains have failed in crystallisation trials. Taken together, these data suggest that the eukaryotic pHAT domains may form homodimers, in agreement with the homodimerisation observed in the hOGA catalytic core crystal structures.

### Eukaryotic pHAT domains are unable to bind acetyl-CoA, but retain an acceptor binding cleft

Prior studies reported that the hOGA pHAT domain is unable to bind acetyl-CoA (Rao, Schüttelkopf *et al*., 2013). To investigate potential species-specific differences, we used surface plasmon resonance to analyse the acetyl-CoA binding properties of OGA pHAT domains from a range of organisms across evolution (**Fig. 1C**). Consistent with earlier reports, none of the eukaryotic pHAT domains tested, including the *Ta*pHAT domain, showed acetyl-CoA binding, while the bacterial HAT from *Oceanicola granulosus* (*Og*HAT) (Rao, Schuttelkopf *et al*., 2013), used here as a positive control, bound acetyl-CoA with a *K*_d_ = 5 μM (**Fig. 1C**). To understand the lack of acetyl-CoA binding, we took advantage of the *Tetrahymena* HAT domain (tGCN5) structure (PDB 1Q2D (Poux & Marmorstein, 2003)) crystallised in a ternary complex with acetyl-CoA and a 19-residue p53-derived acceptor peptide bound in a conserved cleft (Poux & Marmorstein, 2003). Inspection of the superimposed *Ta*pHAT with tGCN5 (RMSD of 4.8 Å across 80 C*α* atoms), reveals steric clashes of the acetyl-CoA with the *Ta*pHAT p-loop, which is crucial in the GCN5 family members for stabilising the negatively charged acetyl-CoA pyrophosphate (**Fig. 1D**) (Dyda *et al*., 2000). Furthermore, the presence of four additional residues in the *Ta*pHAT *α*5 helix is likely to obstruct acetyl-CoA binding (**Fig. 1D and Supplementary Fig. 1D**). This architecture is conserved in the human pHAT domain (**Supplementary Fig. 1D**).

Although acetyl-CoA binding to the pHAT domains may be affected, a comparison of *Ta*pHAT and tGCN5 structures reveals the presence of a partially conserved peptide acceptor binding cleft (**Figs. 1D,E**). This cleft, formed by the *α*1 and the *α*6 helices, is blocked on one side by the presence of additional *α*3 and *α*4 helices in the pHAT domains, which are not present in the canonical GCN5 fold (**Fig. 1E**) (Clements *et al*., 2003). A deeper analysis using the DALI server (Holm, 2020) identified the cryo-EM structure of a macromolecular complex (PDB 7VVU, (Qu *et al*., 2022)) between the yeast nucleosome and a histone acetyltransferase (NuA4) complex where the HAT domain is structurally similar to the *Ta*pHAT domain (RMSD of 3.8 Å across 96 C*α* atoms). An isolated view of this macromolecular complex shows the NuA4 HAT interaction with the histone 4 (H4) tail that penetrates one side of the NuA4 HAT cleft poised for acetylation, and the Polycomb complex related protein Epl1 that interacts on the other side of the cleft and is believed to stabilise NuA4 binding to the nucleosome (**Supplementary Fig. 1E**) (Qu *et al*., 2022). The superposition of the *Ta*pHAT domain structure with this complex reveals that the extra element formed by the *α*3 and *α*4 helices clashes with the H4 tail similarly to the clashes observed with the tGCN5 peptide substrate (**Fig. 1E and Supplementary Fig. 1E**). However, there are no apparent barriers for a putative interaction of the pHAT domain with the Polycomb related protein Epl1 via the pHAT conserved binding cleft (**Fig. 1E and Supplementary Fig. 1F**). Taken together, these data suggest that while the pHAT domains of OGAs cannot bind acetyl-CoA and therefore not possess acetyltransferase activity, it is likely that they retain the ability to interact with other proteins via the remnants of a GCN5 family acceptor binding cleft.

### TaOGA possesses a conserved homodimer interface with disordered pHAT domains

We next performed cryo-EM structure analysis of full-length *Ta*OGA to determine how the pHAT domains are positioned relative to the catalytic domain. An *E. coli* purified sample was vitrified on a grid and imaged by cryo-EM, followed by processing in CryoSPARC (**Supplementary Fig. 3**) (Punjani *et al*., 2017). Particles were extracted using a 480 px box size to locate both the catalytic core and pHAT domains, and initial 2D classification revealed that the homodimeric catalytic core of *Ta*OGA was well-defined, but with no clear density corresponding to the pHAT domains (**Fig. 2A**). 3D reconstruction led to a map that defines the *Ta*OGA homodimer catalytic and stalk domains at 3.0 Å resolution, and an AlphaFold3 (Abramson *et al*., 2024) model was used to guide model building (**Fig. 2B**).

**Figure 2.**
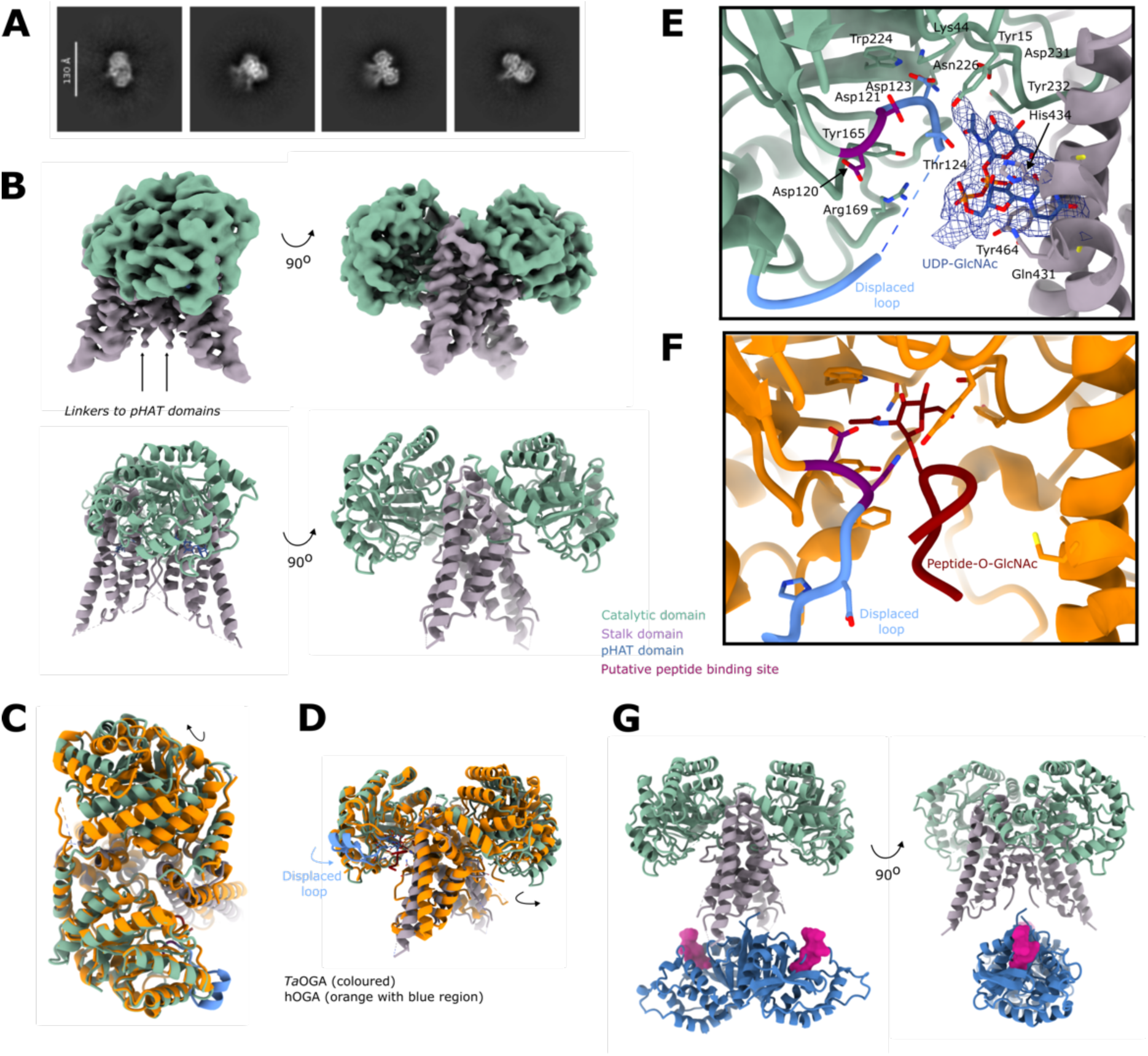
Cryo-EM structural analysis of *Ta*OGA. **A)** 2D classes reveal the catalytic core but the pHAT domains are not visible. **B)** Particles were reconstructed to 3 Å applying two-fold symmetry, without density for the pHAT domains and a density break in the linker region indicating flexibility of the pHAT domains. **C) and D).** Superposition of *Ta*OGA and hOGA with a glycopeptide bound in the active site (PDB 5UN8 (Li, Li, Lu *et al*., 2017)) (orange) shows a conserved dimer interface with tightly connected stalk domain and a looser arrangement between the catalytic domains **(C)** and a displaced loop region in blue **(D)**. Arrows indicate displacement of the catalytic domain of *Ta*OGA relative to the hOGA crystal structure. **E)** Zoom-in of the conserved active site, where the displaced loop occupies the GlcNAc binding site, and non-conserved putative binding site for UDP-GlcNAc. The Asp-Asp catalytic dyad is coloured magenta. **F)** Zoom-in of the active site of hOGA with a glycopeptide. The Asp-Asp catalytic dyad is coloured magenta. **G)** Hybrid model of multi-domain *Ta*OGA using the crystal structure of the *Ta*pHAT domain dimer, the *Ta*OGA cryo-EM structure guided by an AlphaFold3 model, also showing the equivalent position of the P53 peptide obtained from a *Tetrahymena* HAT (PDB 1Q2D (Poux & Marmorstein, 2003)) structure to indicated the position of the pHAT domain putative acceptor peptide binding site (pink).

The dimerisation interface of *Ta*OGA buries 2,700 Å^2^ of the total 20,000 Å^2^ surface area, accounting for 14%, as calculated by the PISA server (Krissinel & Henrick, 2007), which is 20% smaller than the reported for the hOGA crystal structures (Roth *et al*., 2017; Li, Li, Hu *et al*., 2017). The interface is conserved and forms a solvent-exposed hollow channel through the homodimer, with Gln234 (Gln288 in hOGA) from the stalk domains extending into this space and partially narrowing it to pairwise inter-Cα distance 7.2 Å in *Ta*OGA and 8 Å in hOGA **(Fig. 2C, Supplementary Fig. 2)**. The helical region before the linker to the pHAT domain further stabilises the dimer by closing the channel between the monomers. Beyond this point, the linkers are undefined and do not interact with the stalk domains. The interface is looser at the top, between the catalytic domains, and tightens near the active site, suggesting a balance between conformational flexibility while ensuring the active site remains rigid enough for precise substrate positioning. The top of the stalk domains of the *Ta*OGA cryo-EM structure are in the same tight arrangement as in the hOGA catalytic core crystal structures, whereas one of the *Ta*OGA catalytic domains rotates away from the neighbouring stalk domain, which is reflected in a pairwise inter-Cα distance of 33 Å in *Ta*OGA (Lys59) and 40 Å in hOGA (Val113) (**Fig. 2C**).

The substrate binding residues in the active site are conserved and align with the hOGA crystal structure (PDB 5VVO, (Li, Li, Hu *et al*., 2017)), except for a displacement of the loop containing the catalytic acid Asp121 (Asp175 in hOGA) (**Fig. 2E,F**). In the hOGA crystal structure, this loop has a small helical region interacting with the surface of the catalytic core, whereas this region is an unstructured displaced loop showing poor density in the *Ta*OGA cryo-EM structure (**Figs. 2D-F**). A break in the loop density places Asp123 in the GlcNAc binding site, while non-conserved residues Arg169, Gln431, His434, and Tyr464 (corresponding to Phe223, Gly619, Met622, and Ser652 in hOGA, **Supplementary Fig. 2**) coordinate a large additional density **(Fig. 2E)**. While the cryo-EM map suggests that this density could represent a UDP-GlcNAc molecule (a hexosamine biosynthetic pathway product, and the substrate of OGT), with phosphate groups interacting with positively charged residues and π-stacking stabilizing the purine ring, we were unable to confirm either binding to or inhibition of *Ta*OGA. These interactions provide insight into the conserved active site architecture and a non-conserved entry pathway, suggesting potential regulatory mechanisms of *Ta*OGA.

The absence of the pHAT domain from 2D classifications and 3D reconstruction suggests significant flexibility between the catalytic core and pHAT domains, despite the presence of a short linker (STDEYEESTLKNS) connecting these (**Figs. 1A, 2A,B, Supplementary Fig. 2)**. This flexibility is consistent with AlphaFold3 analysis of *Ta*OGA that predicts low pLDDT values for this linker, decoupling the pHAT domains from the catalytic core. Notably, several negative charges in the linker may contribute to electrostatic repulsion of these regions in the homodimer. However, given that both the pHAT and catalytic domains are symmetric homodimers and guided by AlphaFold3 that also predicts a symmetric multi-domain homodimer, we exploited and aligned the two-fold axes in the *Ta*OGA pHAT homodimer crystal structure and the *Ta*OGA homodimer cryo-EM structure to predict the position of the pHAT domains relative to the catalytic cores (**Fig. 2G**). In this model, the pHAT domains are positioned between the stalk domains but do not interact with them. This results in the putative pHAT peptide binding sites pointing away from steric hindrance, facing the glycoside hydrolase active sites. Nevertheless, cryo-EM data suggests potential flexibility in their arrangement relative to each other and/or the catalytic core dimer. Taken together these data suggest that *Ta*OGA, like hOGA, is a homodimer with dimerisation interfaces for the catalytic cores and pHAT domains, although the latter are averaged out due to the flexible linkers under conditions of the cryo-EM experiment.

### hOGA cryo-EM structures suggest discretely disordered pHAT domains

The role of the pHAT domain in regulating hOGA catalytic activity is poorly understood. We next exploited a recently published cell system where endogenous OGT and OGA are both fluorescently tagged, allowing the determination of the effects of exogenous OGA or OGT (or variants thereof) expression on O-GlcNAc homeostasis (Yuan *et al*., 2025; Authier *et al*., 2023). In this assay, expression of a pHAT-less version of hOGA leads to a significant disruption of O-GlcNAc homeostasis compared to a full-length hOGA control, implying an increase of hOGA activity as a function of pHAT domain deletion **(Supplementary Fig. 5)**. However, overexpression of this pHAT-less hOGA harbouring a catalytic inactivation D175N missense mutation prevents the disruption. To further investigate this, we determined the cryo-EM structure of multi-domain hOGA. A construct was designed that lacks the large, disordered loop (residues 397-535, **Fig. 1A**) but retains the pHAT domains. Similar to the *Ta*OGA sample, this hOGA was expressed and purified from *E. coli*, imaged by cryo-EM, and processed using CryoSPARC (**Supplementary Fig. 4**) (Punjani *et al*., 2017). Both 2D and 3D classifications showed particle classes with well-defined catalytic and stalk domains, but none of these included fully defined pHAT domains (**Figs. 3A,B**). Instead, these domains appeared as blurred density in 2D and as an elongated asymmetric density between the two stalk domains in 3D, suggesting that the pHAT domains adopt a range of different angular positions relative to the catalytic core. Less populated 2D classes show pHAT domains in various positions, suggesting pHAT domain positional heterogeneity. To explore this heterogeneity, we next sub-classified the particles to isolate distinct conformational states. This analysis revealed four major conformations, termed I, II, III and IV, each separately subjected to 3D reconstruction and refined to global resolutions ranging from 3.3 to 4.0 Å (**Fig. 3C-F**). An AlphaFold3 model of hOGA was used to guide model building, and the final models across all conformations displayed a near two-fold symmetric catalytic core, forming two glycoside hydrolase active sites composed of the catalytic domain of one monomer and the stalk domain of the other **(Fig. 3B)**. Root-mean-square deviation (RMSD) values of 0.7-0.8 Å confirm that the catalytic core closely resembles that of the published hOGA catalytic core crystal structure (PDB 5VVO, (Li, Li, Hu *et al*., 2017)).

**Figure 3.**
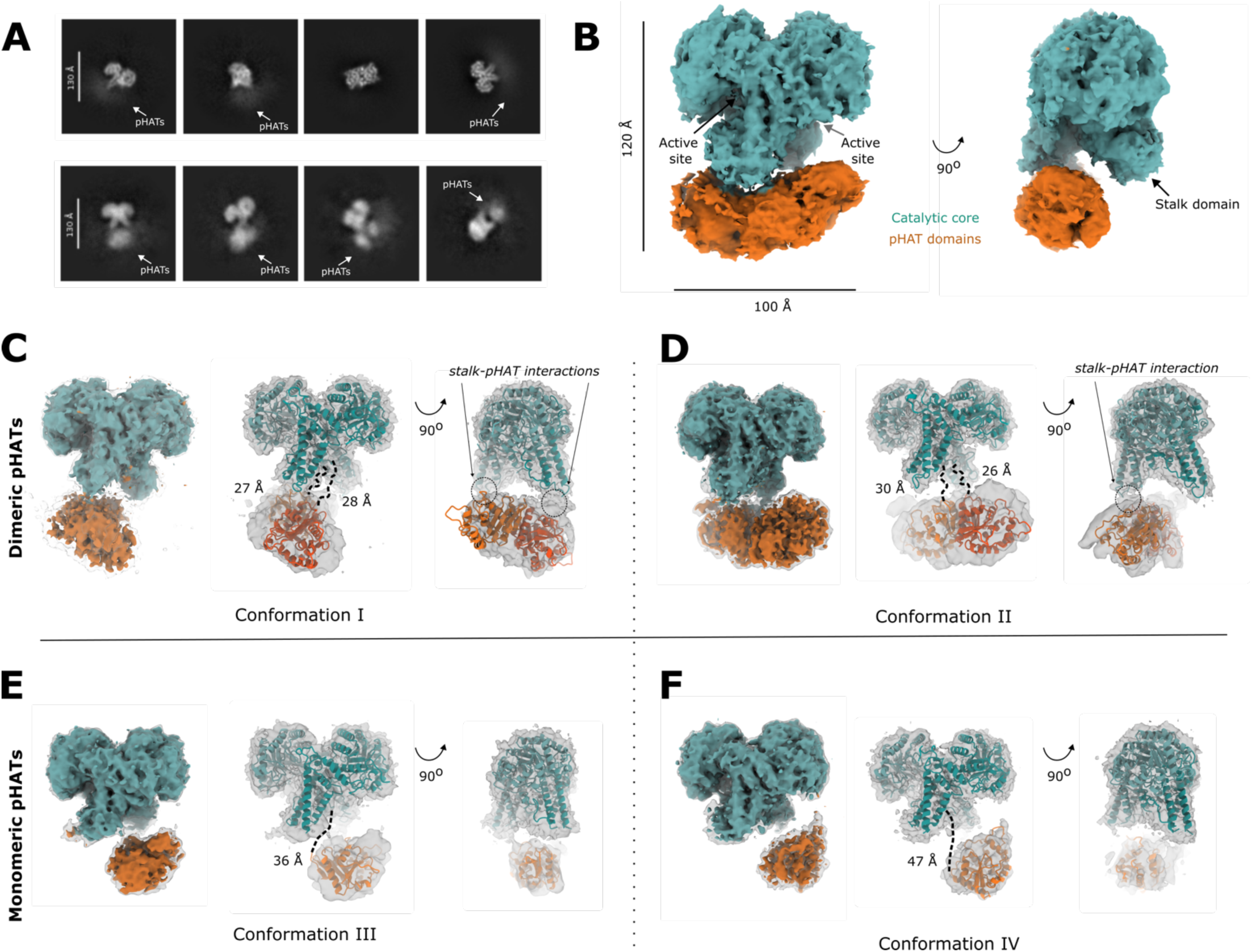
Conformational variability of the hOGA pHAT domains. **A)** 2D classes with 5-6,000 particles reveal a well-ordered catalytic core and blurred density indicates positional heterogeneity for the pHAT domains (upper panel). 2D classes with 1,000 particles reveal pHAT domains in various positions (lower panel). **B)** 140,000 particles were reconstructed to 3 Å. The catalytic core is well-defined, but the pHAT domains appear as an elongated density with multiple positions. Subclassification resolved four different conformations (I – IV) of the pHAT domains. **C) and D)** In conformations I **(C)** and II **(D)** the pHAT domains interact as dimers where the pHAT domains interact with the extended loop of the stalk domains. **E) and F)** In Conformation III **(E)** and IV **(F)** the pHAT domain is found as two separate monomers released from interacting with the stalk domains with only a single pHAT domain being resolved.

While the catalytic core was well-defined, the pHAT domains were not consistently defined in the maps, likely due to their conformational variability. To approximate the positions of the pHAT domains, a blurred density map was generated, providing a coarse view of their locations (**Figs. 3C-F**). A composite map was constructed by combining the high-resolution density of the catalytic core with the blurred density of the pHAT domains, offering an overall structural context. The pHAT domains were subsequently docked into the density using ChimeraX (Meng *et al*., 2023), guided by either a dimeric or monomeric hOGA pHAT model predicted by AlphaFold3.

The four conformations identified represent distinct arrangements of the pHAT domains relative to the catalytic core, highlighting the structural flexibility of hOGA. In conformation I, the pHAT domains form a dimer that interacts with an extended loop region in the stalk domains (residues 594-600**, Supplementary Fig. 2**, **Fig. 4C**). In this state, the density indicates that the dimeric pHAT is bound to the stalk domains, although residual flexibility is implied by the density. Mutating two lysines (Lys597Ala and Lys599Ala) in this loop region in an attempt to decouple this stalk-pHAT interaction does not alter hOGA activity, indicating that this interaction is not critical for regulating the activity of OGA or substrate selection **(Supplementary Fig. 5)**. In conformation II, the pHAT domains retain a dimeric configuration but with only a single pHAT domain interacting with one of the stalk domains (**Fig. 3D**). In this state, the other pHAT domain is stabilized solely through its dimerisation interface, while its connection to the stalk is not defined. Conformation III reveals a monomeric pHAT domain interacting with a single stalk domain, while the other pHAT domain is unresolved (**Fig. 3E**). The linker region connecting the catalytic core to the pHAT domain exhibits some density, indicating partial stabilisation. Indeed, when two key phenylalanines in this linker are mutated (Phe703Ala and Phe704Gly), this leads to reduction in overall hOGA activity in a cell-based system, highlighting the importance of the composition of the linker to retain OGA stability **(Supplementary Fig. 5)**. Conformation IV, similar to conformation III, also features a single monomeric pHAT domain interacting with a stalk domain (**Fig. 3F**). However, in this state, the pHAT domain is rotated closer to the catalytic core and the density for the linker region disappears, perhaps reflecting a loss of ordered interactions with the stalk domain.

**Fig. 4:**
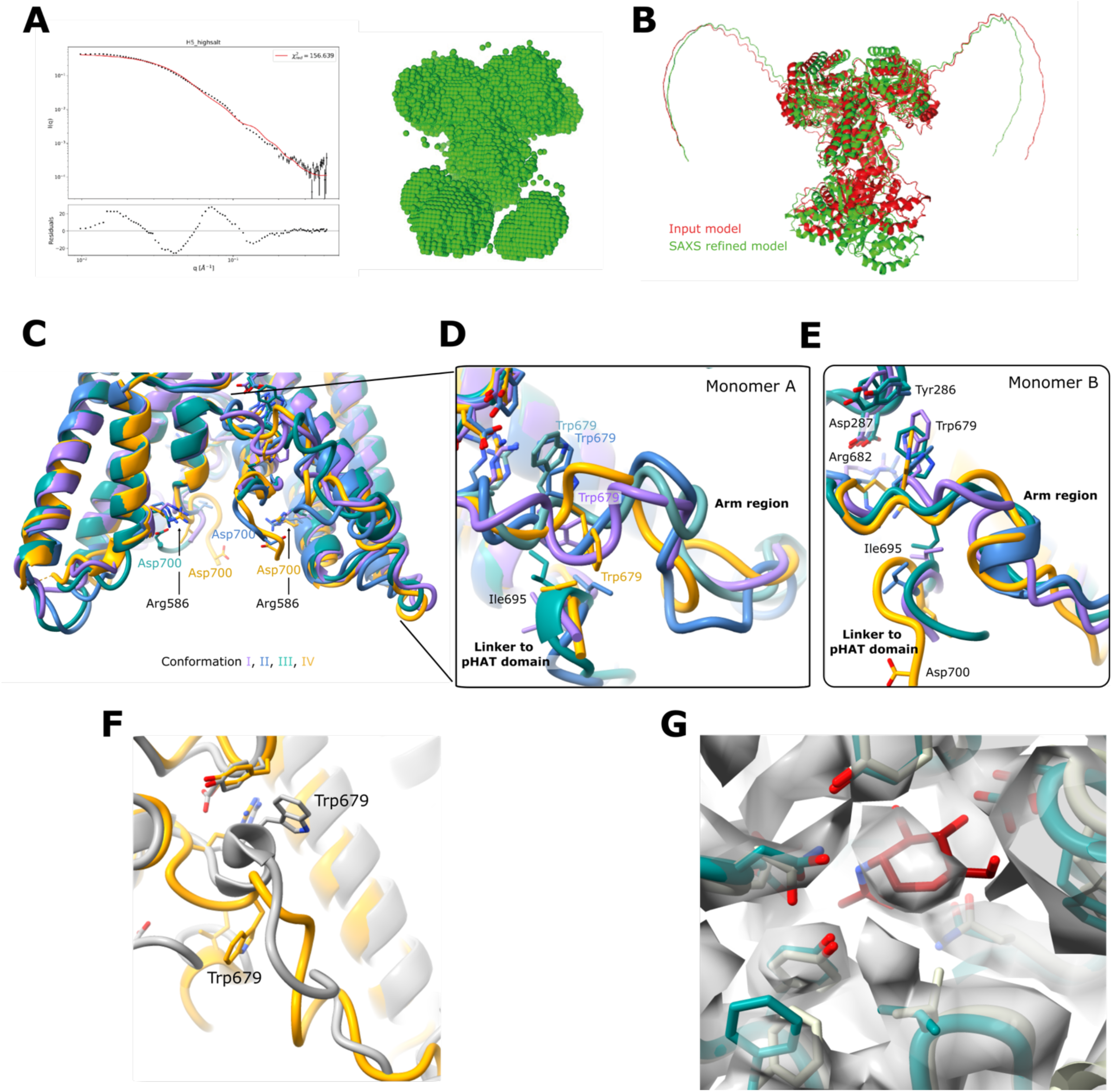
SAXS solution shape of hOGA and pHAT domain-induced conformational changes. **A)** Best fit to the SAXS data and the corresponding model generated for the cryo-EM map of hOGA. **B)** Comparison of the best hOGA SAXS fitted model (green) with the hOGA input model (red) aligned on the stalk region. **C-G)** Structural comparisons of conformations I-IV, covering the stalk domains with visible linkers to the pHAT domains **(C)**, and the arm region from both monomer A **(D)** and monomer B **(E)**. **F)** Structural comparison of the arm region of conformation IV and hOGA crystal structure (gray) (PDB 5VVO (Li, Li, Hu *et al*., 2017)) **G)** Close-up of the active site showing an averaged map of particles from conformation I-IV (teal), with extra density likely corresponding to a GlcNAc molecule (orange sticks).

A comparative analysis of these conformations reveals several trends. Conformation I and II, featuring dimeric pHAT domains, exhibit stronger interactions with the stalk domains, particularly at their loop regions. In contrast, conformations III and IV, characterized by monomeric pHAT domains, exhibit better resolved density for the pHAT domains and reduced variability in their positions. This suggests that dimeric pHAT domains, when stabilized by the stalk domains, adopt more variable conformations, while monomeric domains are more ordered – at least one of them. The four conformations likely represent the four most stable states within a broad conformational landscape of hOGA. Such structural plasticity may be crucial for the function of hOGA, enabling it to interact with a diverse array of substrates or regulatory partners.

### Multi-domain hOGA adopts asymmetric conformations in solution

To further investigate the conformational heterogeneity in the hOGA dimer, we conducted a small angle X-ray scattering (SAXS) experiment using the same construct as used in the cryo-EM experiments. The intensity *I*(*q*) versus the scattering vector modulus (*q*) plots for hOGA are shown in **Supplementary Fig. 6** and further used for a Guinier analysis. The fit of the Guinier residual plot (ln(*I*(*q*)) versus *q*^2^) is acceptable, suggesting that most of the sample is properly folded in solution with little aggregation. The zero-angle scattering intensity values of *I*(0) and the radius of gyration *R*_g_ are given in **Table 2**. These values were used to generate the dimensionless Kratky plots ((*R_g_ q*)^2^ *I*(*q*)/*I*(0) versus (*R_g_ q*)) (**Supplementary Fig. 6**). These data fail to reach a plateau at high *q*, indicating the presence of some flexible parts in the hOGA structure. To gain further insights into the size and shape of the proteins, the pair distance distribution function *p*(*r*) was generated using either the GNOM program of the ATSAS package (Manalastas-Cantos *et al*., 2021) or BIFT (Vestergaard & Hansen, 2006) (**Supplementary Fig. 6**). The GNOM-based *p*(*r*) function goes to zero at 162 Å for hOGA suggesting that these values represent the maximum diameter of the structures. In contrast the BIFT *p*(*r*) function displays a long tail with low amplitude, reaching a diameter of 313 Å for hOGA, evidence for some degree of oligomerisation. The radius of gyration calculated by GNOM aligns well with that from the Guinier fit, indicating consistency between these methods. However, the larger estimation of the radius of gyration by BIFT can be attributed to the influence of the long tail in the *p(r)* functions. This contrast underscores that GNOM primarily captures the compactness of the primary structure, while BIFT may reflect additional contributions from flexibility or transient oligomerization in the sample.

**Table 2:** SAXS data summary.

|  | H5.RSR (hOGA) |
| --- | --- |
| Guinier $R_g$ [Å] | 54.1 +/- 0.3 |
| Guinier $I(0)$ [cm<sup>-1</sup>] | 0.53 +/- 0.002 |
| M.W. ($V_p$) [kDa] | 320.2 |
| M.W. ($V_c$) [kDa] | 282.7 |
| GNOM $D_{max}$ [Å] | 162.0 |
| GNOM $R_g$ [Å] | 53.4 +/- 0.1 |
| GNOM $I(0)$ [cm<sup>-1</sup>] | 0.52 +/- 0.001 |
| BIFT $D_{max}$ [Å] | 313.0 +/- 8.0 |
| BIFT $R_g$ [Å] | 61.6 +/- 1.4 |
| BIFT $I(0)$ [cm<sup>-1</sup>] | 0.55 +/- 0.004 |

To assess the agreement between the cryo-EM map of hOGA and the corresponding SAXS data, a recently published program (Lytje & Pedersen, 2024) was used to fit these data together by calculating the SAXS intensity from the density map. Although the agreement between computed SAXS profiles using high resolution models as templates and the experimental SAXS profiles is not perfect (**Fig. 4A**), the comparison between the cryo-EM model that gives the better fit with the SAXS data shows that the cryo-EM map represents the structures in solution, accounting for the unobserved flexible and disordered regions. To obtain additional structural insights, SAXS data were modelled using high-resolution models obtained from the hOGA catalytic core crystal structure (Roth *et al*., 2017; Li, Li, Lu *et al*., 2017) and a human pHAT 3D model derived from the *Ta*pHAT crystallographic structure via Modeller (Webb & Sali, 2016). These models were further optimised by rigid-body refinement. The resulting models had reduced chi-square values, χ^2^, of 2.9-6.5 for an average oligomerisation of the dimers with ∼60-70% dimers of dimers with an inter-dimer center-of-mass distance of 60-70 Å. These aggregated dimers are also supported by the masses derived from the Porod and the correlation volumes (Manalastas-Cantos *et al*., 2021) (**Table 2**). The dimer structure that gives the best fit to the SAXS data is shown together with the input model which has close to C2 symmetry (**Fig. 4B**). In this best-fit SAXS model, the pHAT domains maintain contact with the stalk domains but are loosely connected to the rest of the structure. These results align SAXS data with the observed flexibility in the hOGA cryo-EM data, corroborating the conclusion that the pHAT domains in hOGA adopt multiple conformations in solution breaking the expected C2 symmetry and generating asymmetric dimers.

### The human pHAT domains induce conformational changes in the arm region leading to active site

In all four conformations of hOGA **(Figs. 3C-F)**, the cryo-EM density maps show a break in density after residue Leu701 within the linker region connecting the catalytic core and the pHAT domain. The linker sequence (LFFQPPPLTPTSKVY, **Supplementary Fig. 2**) is of medium length and proline-rich, a characteristic known to disrupt secondary structures and enhance flexibility (Theillet et al., 2013; Chen et al., 2013) and could therefore contribute to positional freedom of the pHAT domains. The distance from the last residue in a catalytic core secondary structure element before the disordered linker (Leu692) to the first residue of the pHAT domain (Lys713) varies depending on conformation. In conformations I and II, where the pHAT domains interact with the extended loop of the stalk domain, this distance measures 26–28 Å (**Figs. 3C,D**). In conformation II, where the pHAT domain does not directly interact with the stalk domain, this distance extends to 30 Å (**Fig. 3D**). In conformations III and IV, where the pHAT domains are released from stalk domain interactions, this distance increases to 36 Å and 47 Å, respectively (**Figs. 3E,F**). These findings suggest that stalk domain interactions help stabilise the pHAT domains, while the linker enables their conformational flexibility when released. This dynamic positioning of the pHAT domains may have functional implications for hOGA regulation.

Crystal structures of the hOGA catalytic core do not fully define the arm region and the extended linker in the stalk domain. However, our hOGA cryo-EM structures containing the pHAT domain reveal defined density for both the arm and the extended linker regions (albeit at lower resolution, **Supplementary Fig. 4**), suggesting the presence of the pHAT domains increasing order in these regions.

The linkers to the pHAT domains interact with key amino acids on the inside of the stalk domains, anchoring the arm region in a more stable conformation (**Fig. 4C**). The position of the linker and thereby its interaction with the arm region affects the exposure of amino acids leading to the active site (**Figs. 4D,E**). The arm region can undergo a conformational twist, anchoring Trp679 to the linker or towards the catalytic domain (**Figs. 4D**). Specifically in conformation IV, the linker interaction with several amino acids in the stalk domain, where Ile695 of the linker interacts with Trp679 in the arm region, and Asp700 forms a salt bridge with Arg586 in the stalk domain (**Fig. 4E**). In crystal structures with a visible arm region, Trp679 also points towards the catalytic domain (**Fig. 4F**), and depending on anchoring of Trp679, Arg682 either forms an ionic interaction with Asp287, which could stabilise the active site, or occupies the region where Trp679 was anchored to the linker. In this way, the position of the pHAT domain can allosterically regulate the exposure of amino acids in the environment leading to the active site, in agreement with the increase in O-GlcNAcase activity seen upon loss of the pHAT domains (**Supplementary Fig. 5)**.

The cooperativity between the pHAT and stalk domain may also extend into the active site. Unlike the crystal structures of hOGA without pHAT domains, all four conformations display additional density in the active site **(Fig. 4G**). To identify the bound molecule and enhance the signal, all particles were aligned. Superimposing a crystal structure of hOGA with GlcNAc in the active site suggests a density compatible with GlcNAc, presumably co-purified with the enzyme from *E. coli* lysates.

Taken together, the pHAT domain regulates hOGA activity and can adopt various conformations, with its linker stabilising the arm region to expose different amino acids and create distinct environments near the active site. Given the essential role of the stalk domain in substrate recognition (Li, Li, Lu *et al*., 2017; Hu *et al*., 2023) this new insight emphasises the crucial role of the pHAT domain as an allosteric regulator of hOGA.

## Concluding remarks

All proteins capable of hydrolysing O-GlcNAc belong to the GH84 family. Members of this family often couple O-GlcNAcase activity with other functions, possibly to facilitate proper localisation or regulate activity. This functional diversity arises from the fusion of the GH84 catalytic core with additional domains. For instance, bacterial GH84 proteins are frequently associated with domains involved in mediating protein-protein binding interactions, such as coagulant factors, fibronectin type III domains or cohesion and the dockering domains (Drula *et al*., 2022; UniProt Consortium, 2021; Blum *et al*., 2025). In metazoans, the GH84 catalytic activity can appear as a single domain protein, such as the gene *OGA53* in *T. adhaerens* or the short OGA isoform in human cells, which is critical for early neurodevelopment (Liu *et al*., 2012). However, in most metazoans, the GH84 domain of OGA is fused to a histone acetyltransferase-like domain, implying that the combined functions of these two domains are required for the overall protein activity. This suggests cooperation between the catalytic and C-terminal domains for proper O-GlcNAcase function (Liu *et al*., 2021). However, there is substantial conflicting evidence for the function of these C-terminal HAT-like domains. While HAT activity has been proposed, previous work and the data here show that it cannot bind acetyl-CoA and therefore is considered to be a pseudo-histone acetyltransferase (pHAT) of unknown function, retaining a putative peptide binding site. Possible functions could include facilitating hOGA:protein interactions and/or regulating hOGA activity. In this work, we demonstrate that deletion of the pHAT domain induces a more pronounced disruption in O-GlcNAc feedback homeostasis, suggesting increased catalytic activity or broader substrate promiscuity without the presence of the pHAT domain or a direct role for the pHAT domain in maintaining O-GlcNAc homeostasis.

The regulation of OGA appears to have diverged between humans and *Trichoplax*, reflecting adaptations to their respective cellular environments. Notably, the linker between the catalytic core and the pHAT domain, as well as the extended loop in the stalk domain, are not conserved between the two organisms (and smaller/lacking in the *Trichoplax* enzyme). This structural divergence suggests that *Ta*OGA and hOGA may have evolved distinct regulatory mechanisms suited to their functional needs. hOGA exhibits a more intricate regulation, with the pHAT domain positioned to interact with the catalytic core. This arrangement likely provides a fine-tuned mechanism for modulating activity, possibly by influencing substrate recognition or enzyme dynamics. One possible explanation for this regulatory complexity in humans is the likely more complex roles of O-GlcNAc cycling in cellular signalling, metabolism, and disease pathways compared to the simpler multicellular *Trichoplax* organism.

Conformational flexibility is essential in multimers to facilitate interactions with various partners (Pabon & Camacho, 2017; Whitney *et al*., 2016). Though rare, pronounced asymmetry in homodimers can be functionally significant by limiting the active site availability by or enabling interactions with ligands, such as DNA (Swapna *et al*., 2012). Published data suggest that the pHAT domain is critical for hOGA localisation to the nucleus following DNA damage (Liu *et al*., 2021). Consequently, the flexibility of OGA may be crucial for mediating the interactions between the pHAT domain and as yet unknown components of chromatin. The lack of enzymatic activity and the presence of the conserved peptide binding groove in the pHAT domain suggests that this domain has evolved to have a role in mediating specific protein-protein interactions vital for hOGA function.

The evolutionary link between the pHAT and GH84 catalytic domains raises intriguing questions. Disordered linkers in multi-domain proteins often function to increase the local concentration of interacting domains, thereby facilitating allosteric regulation (Huang *et al*., 2020). In the case of OGA, the pHAT domain may play a key role beyond substrate processing — it could act as a structural scaffold for localization, influence substrate specificity, or modulate enzyme activity through cooperative interactions with the catalytic domain. *OGT* missense variants leading to intellectual disability are associated with a (compensatory) loss of OGA mRNA and protein, leading to the intriguing possibility that the resulting loss of such pHAT domain functions could also contribute to the mechanisms underpinning this disease.

Beyond its role in deglycosylation, targeting the pHAT domain or its interaction with the catalytic core could offer a novel approach for therapeutic intervention. Current inhibitors designed to competitively block the GH84 active site face limitations due to the broad range of OGA substrates. A more targeted approach — disrupting the pHAT-GH84 domain cooperativity — could yield greater specificity while preserving basal catalytic function.

Ultimately, hOGA is part of a complex regulatory network governing homeostasis of O-GlcNAcylation, a modification with widespread implications in cellular signalling, metabolism, and disease. Missense variants in hOGA, including in the pHAT domain, are now beginning to be associated with neurodevelopmental disorders (Authier *et al*., 2023). By elucidating the structural mechanisms underpinning hOGA regulation, we provide a platform for dissection of the role of the pHAT domain in mechanisms of modulating O-GlcNAc homeostasis in such disease contexts and beyond.

## Materials & Methods

### Cloning, expression and purification of the recombinant pHAT domain of TaOGA

The pHAT region of *Ta*OGA was isolated from the whole gene (Srivastava *et al*., 2008; Selvan *et al*., 2015) by PCR. Mouse, chicken, zebrafish, *Drosophila* and human pHAT domains were all obtained as codon-optimised gene blocks from Integrated DNA technologies. The sequences coding for pHAT domains were inserted into a pGEX6P1 vector for expression of proteins with a GST tag containing a PreScission protease cleavage site. For protein expression, the pHAT domain plasmids were transformed into *E. coli* BL21(DE3) pLysS. Cell cultures were grown to an OD_600_ of 0.8 and 300 μM IPTG was added to induce protein expression at 16 °C. The cultures were grown for an additional 16 h and harvested by centrifugation. The cells were resuspended in lysis buffer (25 mM Tris buffer pH 7.5, 150 mM NaCl, and 0.5 mM TCEP containing DNase, protease inhibitor cocktail (1 mM benzamidine, 0.2 mM PMSF and 5 µM leupeptin) and lysozyme) and lysed using French press. Cell debris was pelleted by centrifugation and the supernatant was incubated with glutathione Sepharose beads for 2 h on a roller at 4 °C. After extensive wash, the protein was cleaved off the beads with PreScission protease overnight at 4 °C. The proteins were concentrated and loaded onto a 26/600 Superdex 75 column equilibrated with lysis buffer. Fractions confirmed by SDS-PAGE were pooled and concentrated to 30 mg/mL.

### TaOGA pHAT domain crystallisation and structure solution

10 mg/mL of pure *Ta*OGA pHAT domain protein in 25 mM Tris buffer pH 7.5, 150 mM NaCl, and 0.5 mM TCEP was used to screen for crystals at 20 °C using the sitting-drop vapour diffusion method. Each drop contained 0.6 μL of protein solution with an equal volume of the mother liquor (0.1 M HEPES 7.5, 60 mM sodium potassium tartrate and 27.5% PEG 8000). The crystals belonging to the P1 space group grew after 2 days. Crystals were cryo-protected with 15% (v/v) glycerol in mother liquor and frozen in liquid nitrogen. X-ray datasets were collected at the ID30A beamline of the European Synchrotron Radiation Facility (ESRF, Grenoble, France). Data were processed with XDS (Kabsch, 2010). The *Ta*OGA pHAT domain structure was solved using a *Saccharomyces cerevisiae* HAT structure (PDB 4BMH (He *et al*., 2014b)) as the initial phase donor in a molecular replacement experiment using MOLREP (Vagin *et al*., 2010). Refinement was performed with REFMAC5 (Murshudov *et al*., 1997) and model building with COOT (Emsley & Cowtan, 2004). Data collection statistics are listed in **Table 1**. Figures were generated using The PyMOL Molecular Graphics System, Version 2.5 Schrödinger, LLC.

### Cloning, expression and purification of hOGA and TaOGA

The hOGA protein was produced using a plasmid containing a 6xHis tagged N-terminal region of OGA (11-396) followed by the C-terminal region of the protein (535-end) bridged by a GS linker as previously described by (Li, Li, Lu *et al*., 2017; Li, Li, Hu *et al*., 2017). The plasmid backbone pHEX6P1 is a modified version of pGEX6P1 but with a 6His tag replacing the GST tag. This vector adds a PreScission cleavable N-terminal 6His tag. The previously published full length clone of *TaOGA* was used as a template for PCR followed by restriction cloning. The DNA region encoding residues 5-723 were cloned into pHEX6P1 as a *Bam*HI-*Not*I fragment. The resulting human and *Trichoplax* OGA constructs were transformed into *E. coli* BL21(DE3) pLysS for protein expression. Cell cultures were grown to an OD_600_ of 0.6 and induced with 300 mM IPTG at 16 °C overnight. The cultures were harvested by centrifugation at 4000 rpm for 20 min. Cells were resuspended in lysis buffer (25 mM Tris pH 7.5, 150 mM NaCl and 0.5 mM TCEP) and supplemented with DNase, protease inhibitor cocktail (1 mM benzamidine, 0.2 mM PMSF and 5 µM leupeptin) and lysozyme, followed by lysis using a French press. Cell debris was pelleted by centrifugation and the supernatant was incubated with Ni-NTA beads for 2 h on a roller at 4 °C. Beads were extensively washed with lysis buffer and bound proteins were eluted with 250 mM imidazole, followed by overnight incubation with PreScission protease to remove the 6xHis tag. Samples were concentrated and loaded onto a 16/600 Superdex 200 column equilibrated with 50 mM HEPES pH 7.0, 250 mM NaCl and 0.5 mM TCEP). Fractions were confirmed by SDS-PAGE, pooled, concentrated to 10 mg/mL, flash frozen in liquid nitrogen and stored at -80 °C until further use.

### Surface Plasmon Resonance

SPR measurements were collected using a Biacore 3000 instrument (GE Healthcare). OGA pHAT domains were biotinylated by mixing of the protein with amine-binding biotin (Pierce) in 1:1 molar ratio. Streptavidin was immobilized on a CM5 sensor chip surface by amine coupling. 10 mM HEPES pH 7.4, 150 mM NaCl, was used as a running buffer for immobilisation. The surface was activated by 15 min injection of NHS/EDC followed by injection of SA in 10 mM acetate, pH 4.5 until the required density (approximately 9000 relative units (RU)) was achieved and blocked by 4 min ethanolamine injection at 10 μL/min at 25 °C. Biotinylated OGA pHAT domains were captured on the streptavidin surface at approximately 2500 – 4000 RU in running buffer containing 25 mM Tris pH 7.5, 150 mM NaCl, 1 mM DTT and 0.005% Tween 20. Acetyl-CoA was injected in duplicates at threefold concentration series in a range of concentrations 0.2–166.6 μM. Association was measured for 30 s and dissociation for 1 min. All experiments were run at 50 μL/min at 25 °C. All data were referenced for blocked streptavidin surface and blank injections of buffer. Scrubber 2 (BioLogic Software) was used to process and analyse data. Affinities were calculated using 1:1 equilibrium-binding fit.

### Cryo-EM data collection and processing

3 µL of the sample mix (0.5 mg/mL hOGA or *Ta*OGA) was added to freshly glow-discharged (45 s at 15 mA) grids, which were subsequently vitrified at 4 °C and 99% humidity for 3.5 s with blotting force 0. 0.6/1.0 µm 300 mesh AU grids (AuFlat, Protochips) were used for blotting and subsequent plunge freezing, which was carried out on an Vitrobot MarkIV plunge freezer (Thermo Fischer). Data were collected on a Titan Krios G3i microscope (EMBION Danish National Cryo-EM Facility – Aarhus node) operated at 300 KeV equipped with a BioQuantum energy filter (energy slit width 20 eV) and K3 camera (Gatan). A nominal magnification of 130,000x was used, resulting in a pixel size of 0.647 Å^2^/px with a total dose of 59.9 e^−^/Å^2^. The movies were fractionated into 52 frames (1.15 e^−^/Å^2^ per frame) at a dose rate of ∼18 e^−^/px/s and a 1.4 s exposure time per movie.

### Cryo-EM image processing

The images of *Ta*OGA dataset were processed using the pipeline outlined in **Supplementary Fig. 3**. Micrographs were motion corrected (Rubinstein & Brubaker, 2015) and CTF estimated (Zivanov *et al*., 2020) using CryoSPARC (Punjani *et al*., 2017). Particles were picked using a circular blob and aligned by 2D classification. Small subsets of the 2D classes were selected and used to generate ab initio volumes. Particles were primarily sorted alternating between heterogeneous refinement, with ab-initio volumes created from selected 2D classes, and sorting by creating two ab-initio volumes and carrying on with the most well-defined particle stack. The particles were initially extracted in a bigger box to ensure that the pHAT domains were included. However, after realising the pHAT domains could not be resolved, the particles were re-extracted in a smaller box and finally refined in a non-uniform and a local refinement (Punjani *et al*., 2020).

The images of the hOGA dataset were processed using the pipeline outlined in **Supplementary Fig. 4**. The micrographs were motion corrected (Rubinstein & Brubaker, 2015) and CTF estimated (Zivanov *et al*., 2020) using CryoSPARC (Punjani *et al*., 2017). Particles were picked using a circular blob and aligned by 2D classification. A small subset of the 2D classes were selected and used to reconstruct ab initio volumes. One of the ab initio volumes was used for template picking in all micrographs to extract particles. Particles from selected 2D classes were used as templates for re-extraction in a bigger box. Particles were sorted in 2D to remove junk, and reconstruction in several *ab initio* volumes that sorted dimeric from monomeric particles and junk. The particles were also sorted into four main conformations with these strategies. Finally, by using the “particle sets” tools, we separated particles overlapping in several conformations, to ensure particles in each conformation were unique. Each conformation was refined both in a non-uniform and a local refinement (Punjani *et al*., 2020).

### Model building, refinement and validation

AlphaFold3 models were manually docked in the cryo-EM volumes and fitted to the map with geometry restraints using Namdinator (Kidmose *et al*., 2019). Real-space refinement of the structures was done in Phenix (Liebschner *et al*., 2019), and model building and analysis were performed in Coot (Emsley & Cowtan, 2004) (**Table 3**). Figures for the cryo-EM volumes and models were made in ChimeraX (Meng *et al*., 2023).

**Table 3:**
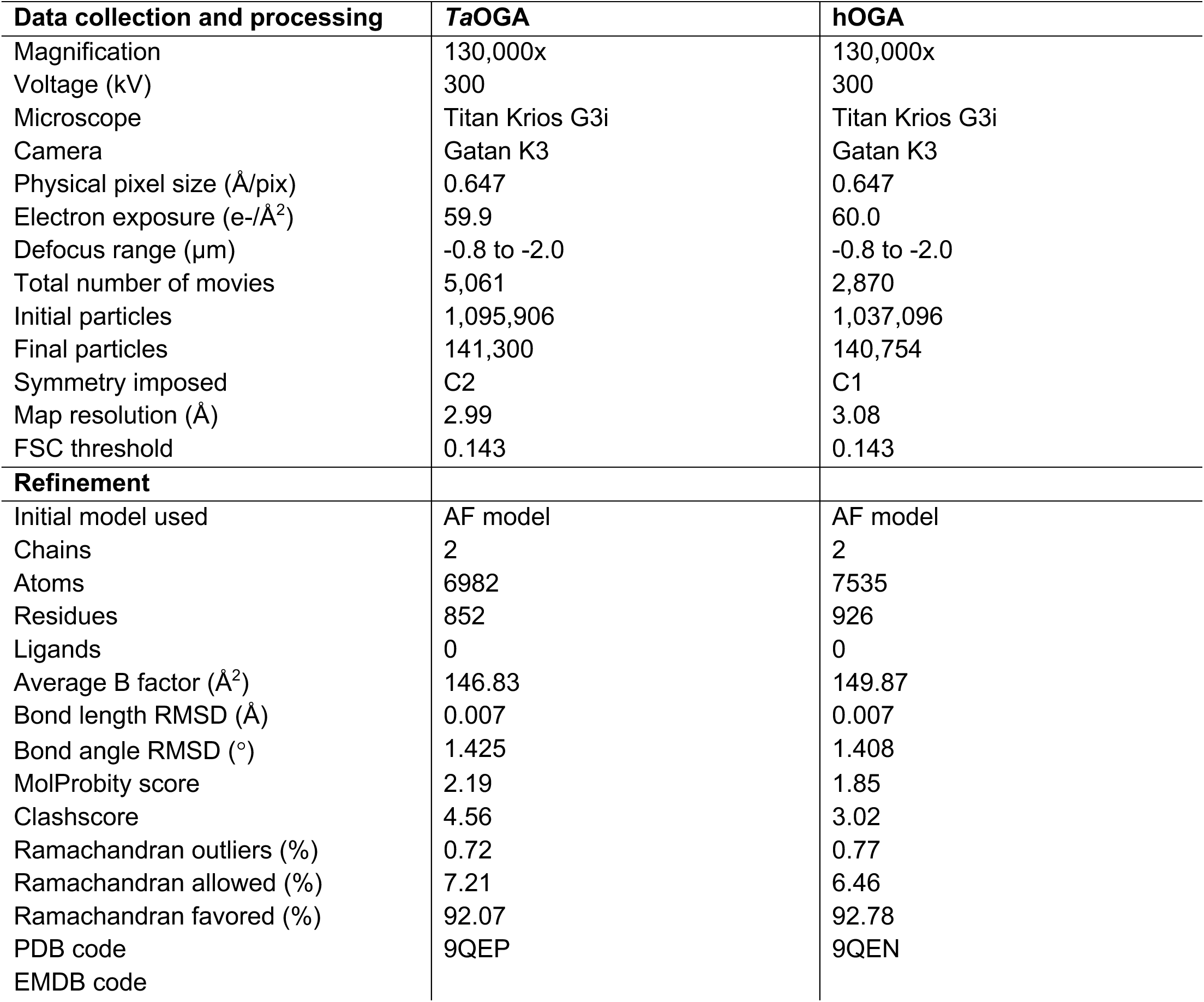
Data collection and refinement statistics for cryo-EM analysis.

### SAXS data collection and analysis

Small-angle X-ray scattering was performed at the in-house facility at Aarhus University (Lyngsø & Pedersen, 2021). It is a modified Bruker AXS NanoSTAR that uses an Excillum Galium liquid metal jet X-ray source and a homebuilt scatterless pinhole (see for example European Patent EP13159569.6 (Pedersen, 2014).), and is equipped with an automated sample handler, which injects the sample into a reusable quartz capillary allowing samples and buffers for background subtraction to measure in the same capillary. Water at 20 °C was used for absolute calibration. The resulting data are shown as the intensity *I*(*q*) versus the modulus of the scattering vector *q* in the plots. Resulting datasets were plotted in a log-log plot, and a Guinier plot of data and linear fit of ln(*I*(*q*)) versus *q*^2^ was performed to obtain values for *I*(0) and the radius of gyration *R*_g_. A dimensionless Kratky plot of (*R_g_ q*)^2^ *I*(*q*)/*I*(0) versus (*R_g_ q*) was made to identify the influence of flexible parts. Indirect Fourier transformations were performed using the GNOM program of the ATSAS package (Manalastas-Cantos *et al*., 2021) and BIFT (Vestergaard & Hansen, 2006). Masses were estimated from the Porod volume *V*_p_ and the correlation volume *V*_c_ as determined by the ATSAS package. The agreement of the cryo-EM map and the SAXS data was checked using a recently published program, which scans a threshold value for the map and creates dummy atom models and determines the reduced chi-squared for each model, thus, obtaining the model from the map that gives the best agreement with the SAXS data (Lytje & Pedersen, 2024). To obtain more information on solution structure of hOGA, the SAXS data were further analysed using high-resolution models of the full-length hOGA obtained via modeller software (Webb & Sali, 2016). The scattering for this structure was calculated and compared to the SAXS data using an home-written program, which was described previously (Bærentsen *et al*., 2023; Vilstrup *et al*., 2020; Steiner *et al*., 2018; Harwood *et al*., 2021). In brief, the program uses an equivalent model approximation for the non-hydrogen atoms with a Gaussian form factor, adds a hydration layer around the protein, and calculates the SAXS intensity on absolute scale using the Debye equation (Debye, 1915). The program allows rigid-body optimization of the structure employing soft connectivity and excluded volume restraints. A structure factor for unspecific oligomers can also be fitted to account for example for presence of a fraction of unspecific dimers (Bærentsen *et al*., 2023).

The hOGA dimer was divided into eight bodies by identifying flexible and compact parts of the structure and considering that the structures should have some degree of freedom to relax during the optimization. The agreement with the SAXS data was optimized by performing refinements of the starting structure using random translations and rotations. For each optimization, 10 runs were performed to investigate variations in the structure and to find the structure where the calculated scattering has the best agreement with the measured SAXS data. A satisfactory fit in terms of reduced chi-square, χ^2^, could not be obtained without including an oligomer structure factor.

### Determination of O-GlcNAc dyshomeostasis using a double fluorescence-labelled cell line

A previously engineered cell line incorporating endogenously labelled OGT-sfGFP/mScarlet3-OGA was used to measure effects on O-GlcNAc homeostasis as a result of transfection of mTagBFP2-tagged exogenous hOGA and variants thereof (Yuan *et al*., 2025; Authier *et al*., 2023) Briefly, 3.5 × 10^5^ cells of this line were transfected with 1.5 µg of DNA plasmid and 3 µL of Lipofectamine 2000 in a 12-well plate setting according to the manufacturer’s instructions. 24 h post-transfection, the medium was changed, and cells were analysed in a flow cytometer (NovoCyte Quanteon 4025) 48 h after transfection. For fluorescence detection, harvested cells were stained with the live/dead dye RedDot^TM^2 (Biotium), and the live, mTagBFP2-positive transfected cells were gated for OGT-sfGFP and mScarlet3-OGA fluorescence detection, and the ratio between these plotted as a measure of induction of O-GlcNAc dyshomeostasis.

## Acknowledgements

This work was funded by a Wellcome Trust Investigator Award (110061), a Novo Nordisk Fonden Laureate award (NNF21OC0065969) and a Villum Fonden Investigator (00054496) to D.M.F.v.A. Supported in part by the Danish Research Institute of Translational Neuroscience – DANDRITE of the Nordic-EMBL Partnership for Molecular Medicine and Lundbeckfonden. We would like to thank ESRF (beamline ID30A) for the synchrotron time. We are also grateful to Iva Hopkins-Navratilova and Tonia Aristotelous from the University of Dundee Drug Discovery Unit for their assistance in performing the SPR experiments. We also extend our gratitude to the FACS Core Facility at Aarhus University for their support.

## Author contributions

S.G.B, and D.M.F.v.A conceived the study; S.G.B., S.B.H., O.G.R., T.D., T.B., J.S.P., K.L. and A.G. performed biochemical experiments and structural biology; A.T.F. performed molecular biology; H.J.Y. performed cellular experiments. S.G.B, S.B.H. and D.M.F.v.A. interpreted the data and wrote the manuscript with input from all authors.

---

**Supplementary Fig. 1:**
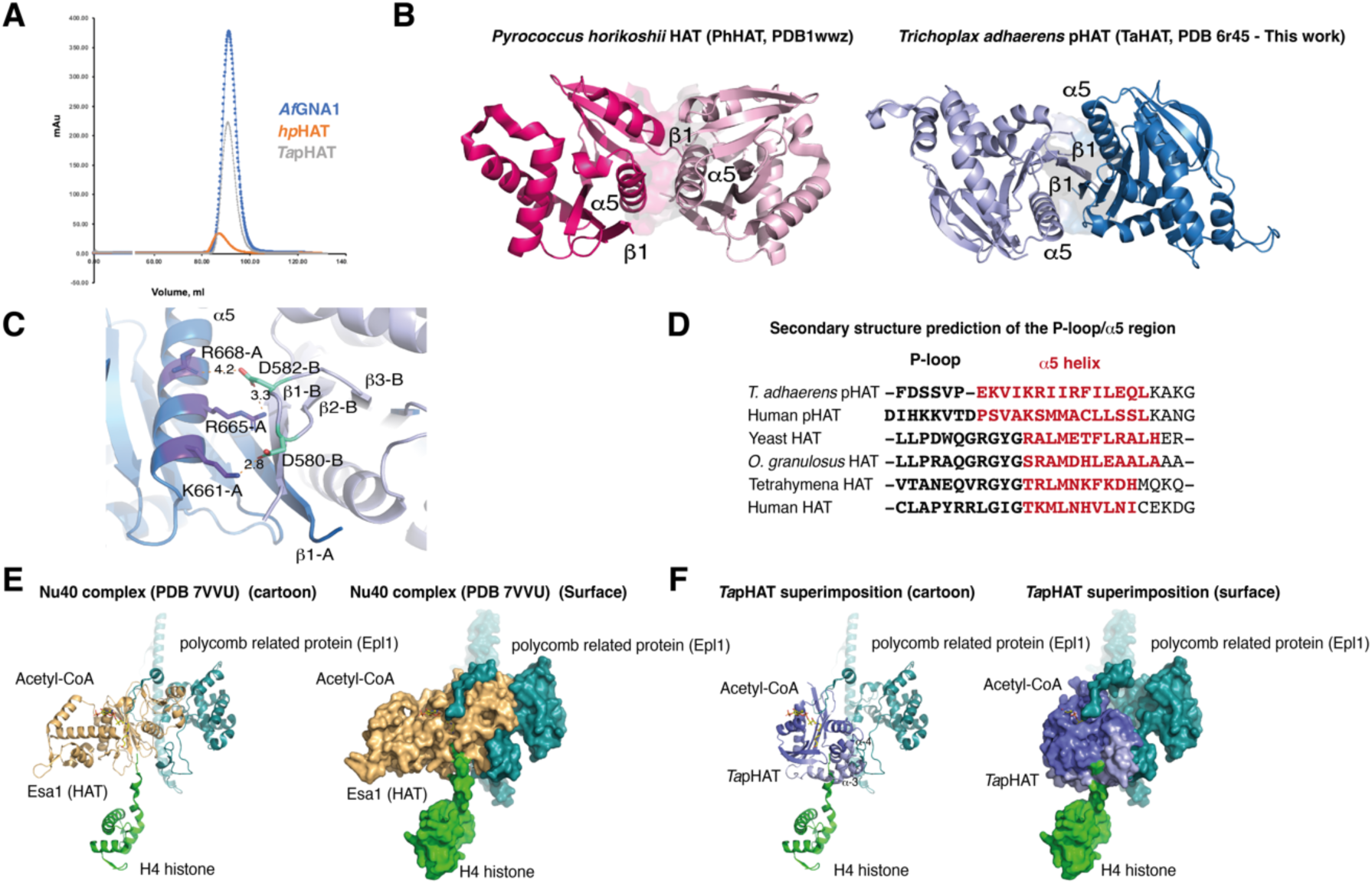
Crystal structure of the *Ta*OGA pHAT domain dimer. **A)** Size exclusion chromatography profile of *Trichoplax* and human pHAT domains using *Af*GNA1 as control. **B)** Dimeric arrangement of the Archaeota HAT domain from *Pyrococcus horikoshii* (Kunishima, N, 2005) (red and pink) and the *Ta*OGA pHAT domain (light blue and blue). **C)** Hydrogen bonds and electrostatic interactions found in the dimerisation interface. Monomer A is coloured marine, and the *α*5 helix-interacting residues are in blue magenta. Monomer B is coloured slate, and the interacting residues from the *β*2-*β*3 loop are in teal. **D)** Sequence alignment of the pHAT domain p-loop and *α*-5 regions across evolution. **E)** Details of the Nu40 HAT complex elements. **F)** Details of the superposition of the *Ta*pHAT domain onto the Nu40 HAT complex, highlighting potential steric clashes between the *Ta*pHAT domain and the H4 histone in this specific arrangement.

**Supplementary Fig. 2:**
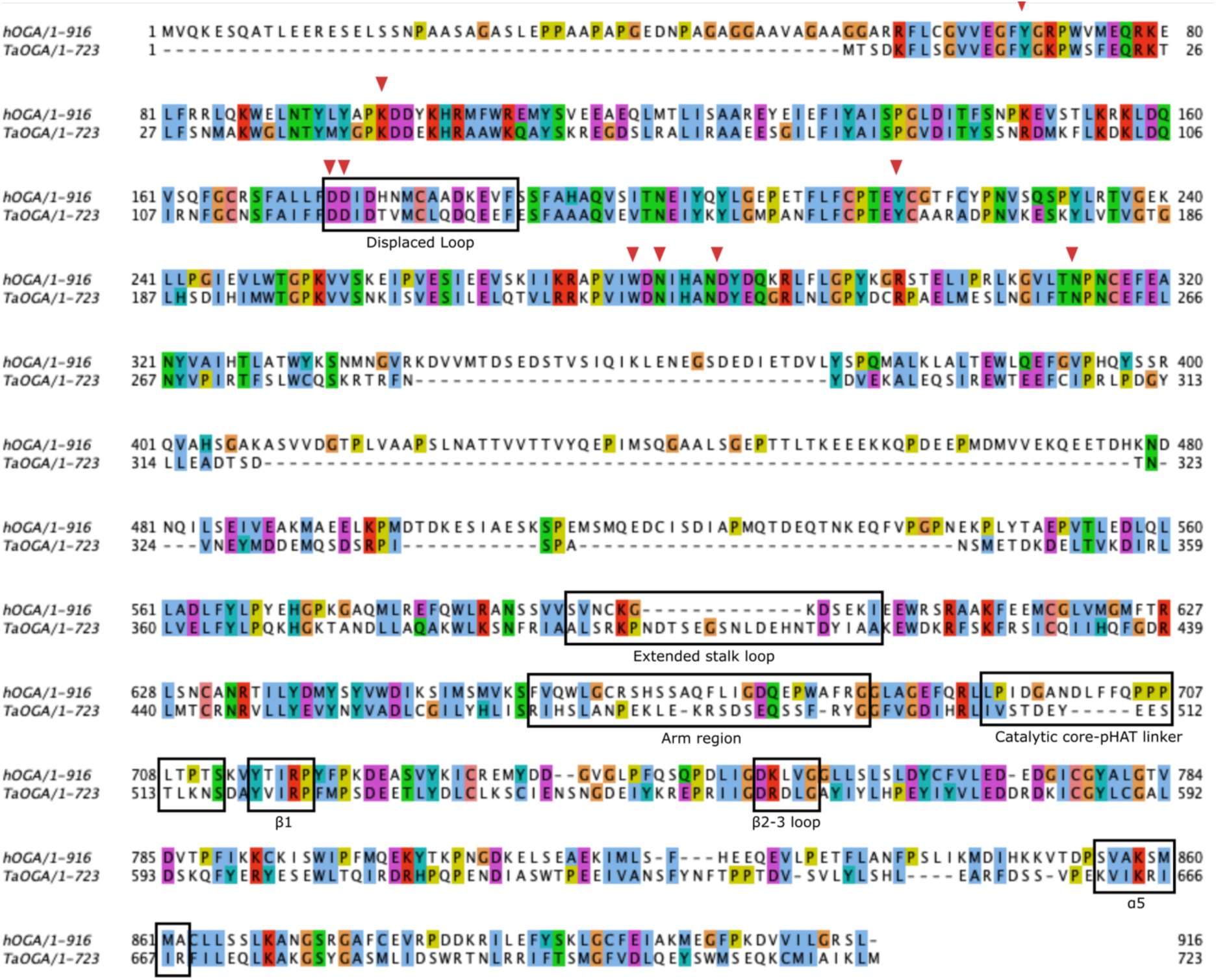
Sequence alignment of *Ta*OGA and hOGA. Red arrows indicate residues coordinating GlcNAc in the active site. Specific regions are highlighted in boxes.

**Supplementary Fig. 3:**
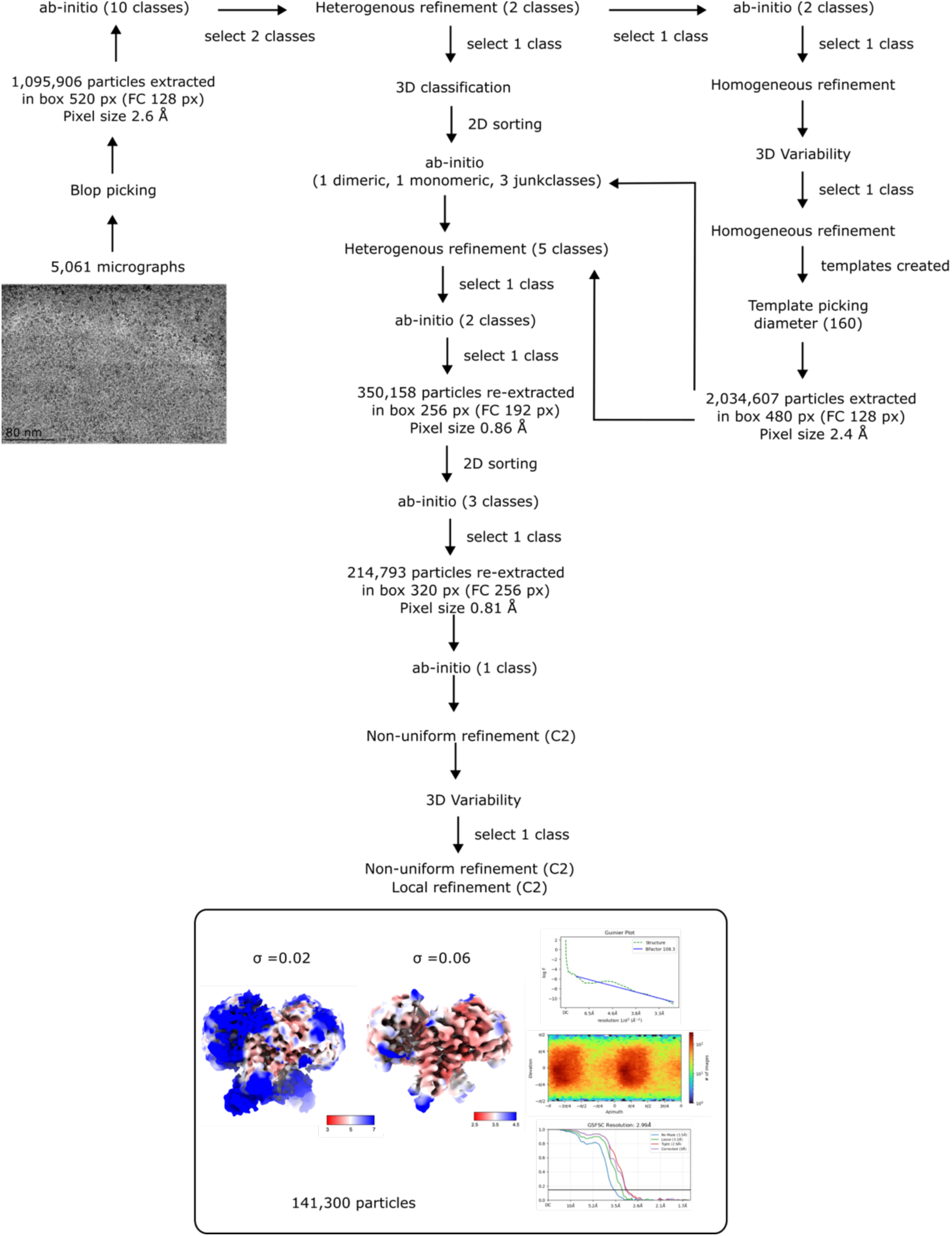
Cryo-EM pipeline for *Ta*OGA.

**Supplementary Fig. 4:**
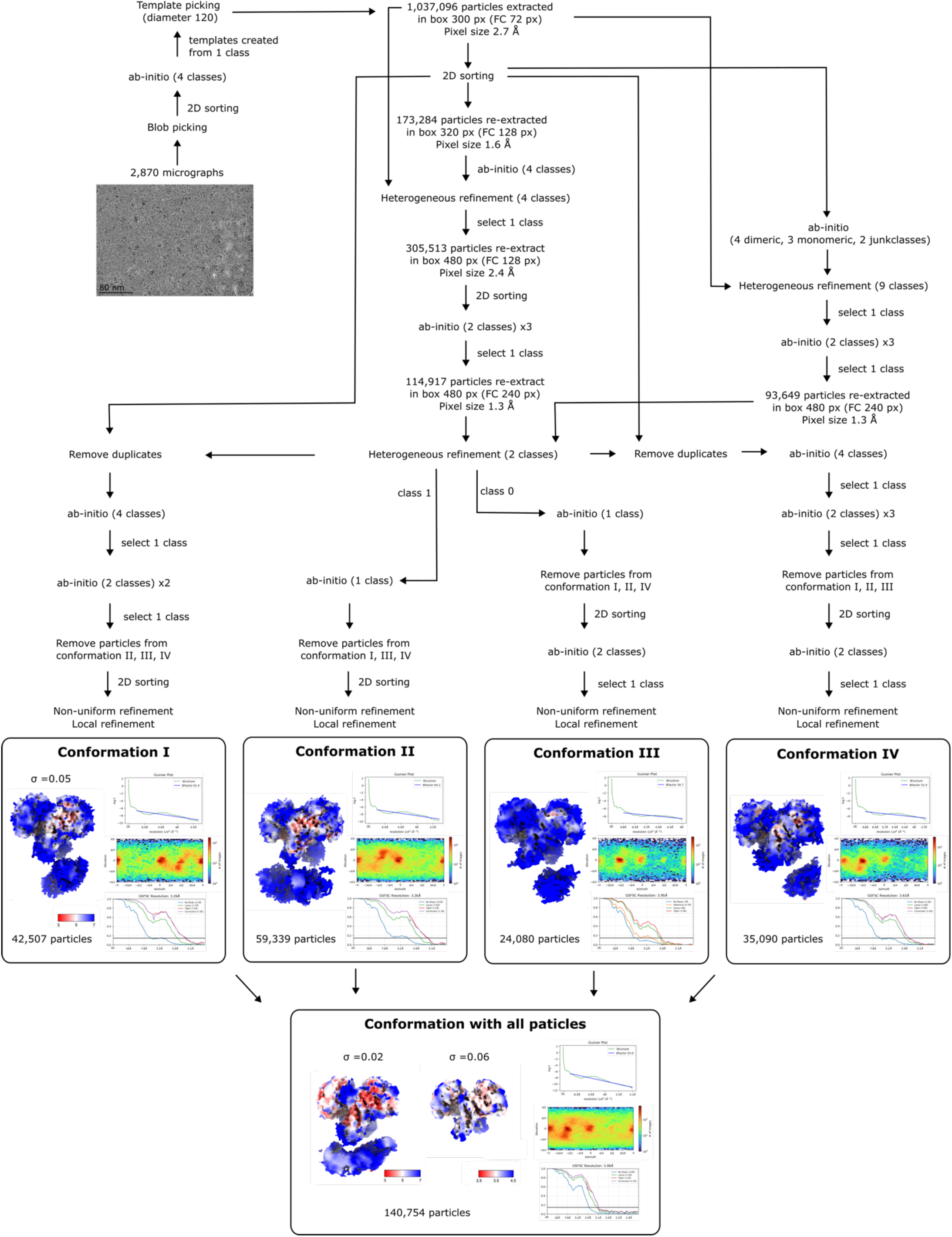
Cryo-EM pipeline for hOGA.

**Supplementary Fig. 5:**
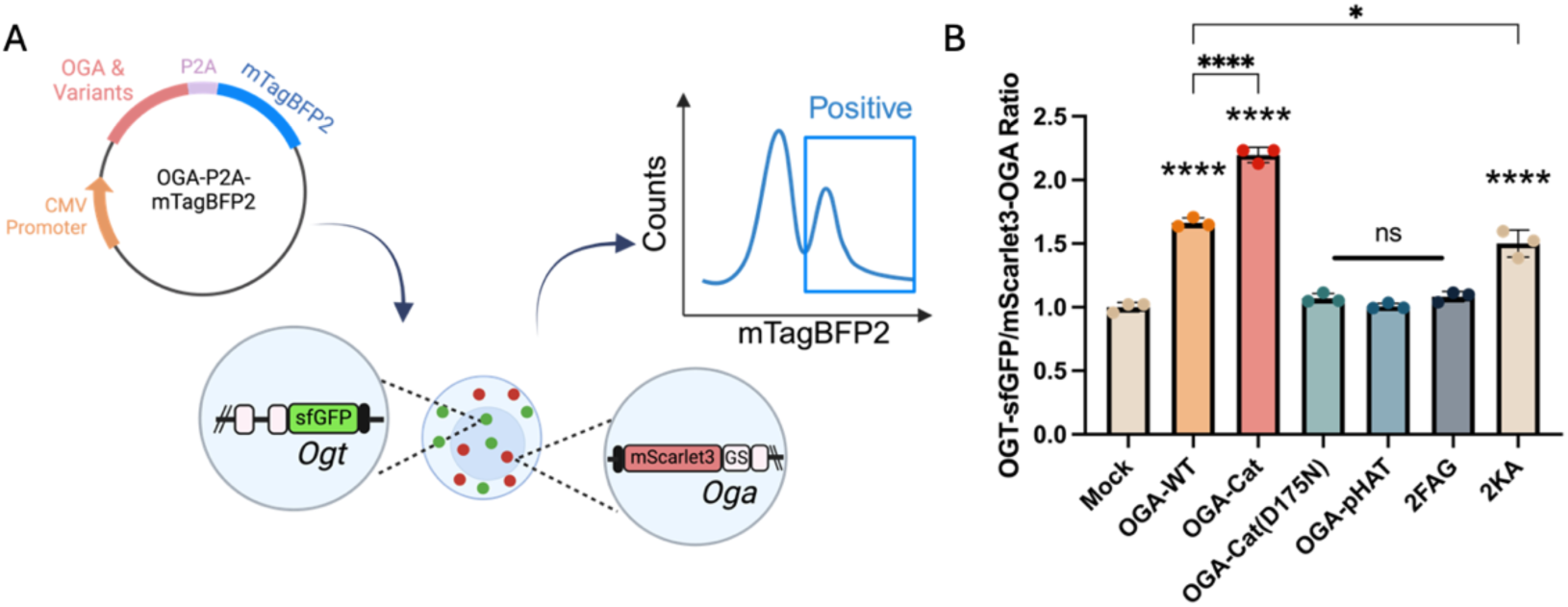
Disruption of O-GlcNAc homeostasis by hOGA variants. **A)** Brief outline of the double-fluorescently labelled mESC system to detect changes in O-GlcNAc homeostasis (as measured by changes in endogenous OGT/OGA levels) as a result of transfecting exogenous mTagBFP2-labelled hOGA variants (see M&M for details) **B)** Ratio of OGT/OGA levels with a range of hOGA variants, including controls. Compared to mTagBFP2 only transfection (Mock), wild type OGA (OGA-WT) transfection increased OGT and decreased OGA endogenous protein levels, resulting in an elevated OGT/OGA ratio. Removal of the pHAT domain (OGA-Cat) led to a more pronounced increase in the OGT/OGA ratio, suggesting an exacerbated disruption in O-GlcNAc homeostasis. However, introducing the D175N inactivating mutation into this construct (OGA-Cat(D175N)) abolished this effect. Transfection with the hOGA pHAT domain alone (OGA-pHAT) induced no changes in OGT/OGA ratio. A hOGA double lysine and double phenylalanine mutant (2KA, 2FAG, as referenced in the main text) are also included. Ordinary one-way ANOVA was performed (n = 3 technical replicates). Statistical significance between the Mock and each variant is directly indicated above the respective groups. Significance for selected pairwise comparisons is displayed with connecting lines, with adjusted *p* values denoted as follows: ns (*p* > 0.05), * (*p* < 0.05), *** (*p* < 0.001), and **** (*p* < 0.0001). Error bars represent the mean ± SD.

**Supplementary Fig. 6:**
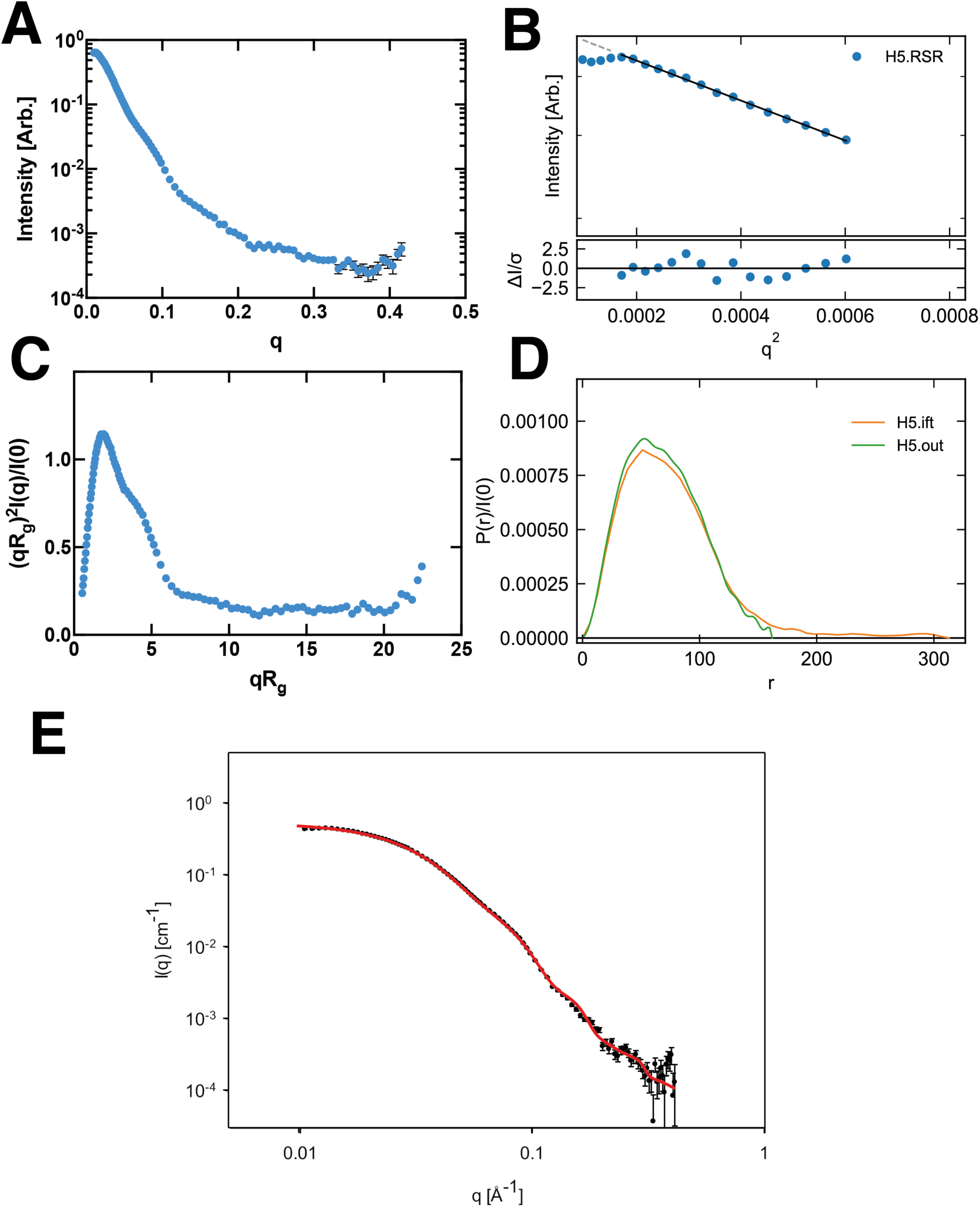
SAXS data summary figure for hOGA. A) Scattering profile on a log-linear scale. B) Guinier fit(s) (top) and fit residuals (bottom). C) Dimensionless Kratky plot. Dashed lines show where a globular system would peak. D) *p(r)* function(s), normalized by *I(*0). E) Fit to the SAXS data for the best models from rigid-body optimization. The lower black points represent hOGA. The red curves are the model fits.

